# Catalytically inactive subgroup VIII receptor-like cytoplasmic kinases regulate the immune-triggered oxidative burst in *Arabidopsis thaliana*

**DOI:** 10.1101/2024.05.30.596543

**Authors:** Márcia Gonçalves Dias, Thakshila Dharmasena, Carmen Gonzalez-Ferrer, Jan Eric Maika, Maria Camila Rodriguez Gallo, Virginia Natali Miguel, Ruoqi Dou, Melissa Bredow, Kristen R Siegel, Richard Glen Uhrig, Rüdiger Simon, Jacqueline Monaghan

**Author notes:** Equal contribution (Márcia Gonçalves Dias, Thakshila Dharmasena).

## Abstract

Protein kinases are key components of multiple cell signaling pathways. Several protein kinases of the receptor-like cytoplasmic kinase (RLCK) family have demonstrated roles in immune and developmental signaling across various plant species, making them a family of interest in the study of phosphorylation-based signal relay. Here, we present our investigation of a subfamily of RLCKs in *Arabidopsis thaliana*. Specifically, we focus on subgroup VIII RLCKs: MAZ and its paralog CARK6, as well as CARK7 and its paralog CARK9. We found that both MAZ and CARK7 associate with the calcium-dependent protein kinase CPK28 *in planta,* and furthermore that CPK28 phosphorylates both MAZ and CARK7 on multiple residues in areas that are known to be critical for protein kinase activation. Genetic analysis suggests redundant roles for MAZ and CARK6 as negative regulators of the immune-triggered oxidative burst. We find evidence that supports homo– and hetero-dimerization between CARK7 and MAZ, which may be a general feature of this protein family. Multiple biochemical experiments suggest that neither MAZ nor CARK7 demonstrate catalytic protein kinase activity *in vitro.* Interestingly, we find that a mutant variant of MAZ incapable of protein kinase activity is able to complement *maz-1* mutants, suggesting noncatalytic roles of MAZ *in planta*. Overall, our study identifies subgroup VIII RLCKs as new players in Arabidopsis immune signaling and highlights the importance of noncatalytic functions of protein kinases.

## Introduction

Phosphorylation is one of the most influential post-translational modifications shaping eukaryotic proteomes. Phosphorylated proteoforms can differ in their activation status, localization, stability, or ability to interact with other proteins, increasing the diversity of functions attributed to the products of single genes. Moreover, phosphorylation events occur in a cell-autonomous manner, which is critical to the ability of individual cells to respond to stress signals quickly. Phosphorylation is catalyzed by protein kinases that hydrolyze the γ-phosphate from ATP and transfer it to substrate proteins, typically on serine, threonine, or tyrosine residues. Compared to the metazoan protein kinase superfamily, the plant protein kinase superfamily is hugely expanded. For example, while there are ∼500 protein kinases encoded in the human genome, there are ∼1,400 protein kinases encoded in the genome of the angiosperm model plant *Arabidopsis thaliana,* and even more in plant genomes with higher ploidy levels. Within the superfamily, there are several subfamilies of protein kinases that differ in their enzymatic and substrate recognition properties, and can be phylogenetically organized based on their unique features (Lehti-Shiu and Shiu, 2012). The largest protein kinase families in plants include transmembrane receptor-like kinases (RLKs) and cytoplasmic kinases of the mitogen-activated protein kinase (MAPK), Ca^2+^-dependent protein kinase (CDPK), and receptor-like cytoplasmic kinase (RLCK) subfamilies. A typical stress signaling pathway includes the activation of an RLK by binding an extracellular ligand followed by the trans-phosphorylation and activation of cytoplasmic protein kinases which then phosphorylate various substrates that result in signal transduction and temporary cellular reprogramming.

As in many other stress pathways, protein kinases are integral to immune signaling. Immune sensing is achieved by pattern recognition receptors (PRRs) in the plasma membrane, many of which are either transmembrane RLKs or non-kinase receptor-like proteins (RLPs) (DeFalco and Zipfel, 2021). Typically, serine/threonine protein kinases have an arginine preceding a conserved aspartate residue within their catalytic loops and are thus referred to as RD kinases (Dardick and Ronald, 2006). Most immune-related PRRs are non-RD kinases that lack the aspartate residue and are less catalytically active than their RD counterparts (Dardick and Ronald, 2006; Tang *et al*., 2017). Due to this, PRRs generally function in large multiprotein immune complexes, and often require association with a co-receptor for signal transduction, such as BRASSINOSTEROID INSENSITIVE 1 ASSOCIATED KINASE 1 (BAK1) (Nam and Li, 2002; Chinchilla *et al*., 2007; Heese *et al*., 2007; Monaghan and Zipfel, 2012). PRRs bind highly conserved microbial features known as microbe-associated molecular patterns (MAMPs) via their extracellular ligand-binding domains. Highly studied MAMPs include the 22 amino acid epitope of the bacterial motor protein flagellin (flg22), the 18 amino acid segment of elongation factor Tu (elf18), or components of bacterial or fungal cell walls (Shu *et al*., 2023). Following immune induction, endogenous small peptides called phytocytokines are released and can bind additional PRRs and amplify and propagate the signal (Segonzac and Monaghan, 2019), together resulting in the activation of other protein kinases, the release of reactive oxygen species (ROS) into the apoplast, an increase in cytoplasmic Ca^2+^, and ultimately, transcriptional reprogramming (DeFalco and Zipfel, 2021).

Involved in growth, development and stress signaling, the 149 RLCKs in Arabidopsis cluster into 17 subgroups referred to as RLCK-II and RLCK-IV – RLCK-XIX (Shiu and Bleecker, 2001; Liang and Zhou, 2018). The most studied RLCKs belong to the 46-member subgroup VII, including BOTRYTIS INDUCED KINASE 1 (BIK1) and closely related PBS1-like (PBL1) proteins (Lu *et al*., 2010; Zhang *et al*., 2010), which form dynamic complexes with multiple PRRs (Liang and Zhou, 2018). Signaling through BIK1 and PBL1 proteins is genetically required for immune responses, and these RLCKs are directly targeted by secreted pathogenic proteins known as effectors to suppress immune responses (Veronese *et al*., 2006; Lu *et al*., 2010; Zhang *et al*., 2010). Although BIK1 and related subgroup VII RLCKs have emerged as key regulators of immunity over the past decade, the earliest reports of RLCKs involved in pathogen defense surrounded the tomato kinases PSEUDOMONAS SYRINGAE PATHOVAR TOMATO (Pto) and FENTHION SENSITIVITY (Fen) (Chandra *et al*., 1996; Jia *et al*., 1997). Both Pto and Fen interact with immune receptor PTO RESISTANCE AND FENTHION SENSITIVITY (Prf) (Mucyn *et al*., 2006, 2009), which belongs to a cytoplasmic immune receptor family known as the nucleotide-binding, leucine-rich repeat (NLR) family. Whereas PRRs detect the presence of microbes in the extracellular space, NLRs detect the presence of pathogens that have actually infected cells (Kourelis, 2023). Activated NLRs directly mount a robust immune response that results in cell death to clear the infection and protect neighbouring tissues. In this pathway, Pto and Fen are not classical ‘signal transducers’; rather, modification to their proteoforms serves as a marker for the presence of the secreted protein effectors AvrPto and AvrPtoB from the virulent bacterial pathogen *Pseudomonas syringae* pv. *tomato* that in turn activates Prf (Mucyn *et al*., 2006, 2009). In addition, several RLCKs have been shown to function as decoy substrates for effectors that result in NLR activation (Wang *et al*., 2015; Martel *et al*., 2020), illustrating the varied and complex roles that RLCKs play in immune signaling.

BIK1 protein accumulation is controlled through an intricate interplay between phosphorylation and ubiquitination, which is thought to optimize immune signaling (Dias *et al*., 2022). Unactivated BIK1 is polyubiquitinated by related E3 ubiquitin ligases PLANT U-BOX 25 (PUB25) and PUB26 (Wang *et al*., 2018*a*). The calcium-dependent protein kinase CPK28 associates with and phosphorylates BIK1, PUB25, and PUB26 (Monaghan *et al*., 2014; Wang *et al*., 2018*a*). Although we are still investigating the function of CPK28-mediated phosphorylation on BIK1, it has been shown that CPK28-mediated phosphorylation on PUB25 and PUB26 results in their activation and subsequent polyubiquitination of BIK1 (Wang *et al*., 2018*a*). We recently showed that CPK28 can also phosphorylate the Raf-like kinases MRK1, RAF26, and RAF39, which function in stomatal immunity (Dias *et al*., 2023). Here, we focus on the subgroup VIII RLCKs MAZZA (MAZ) and CYTOSOLIC ABA RECEPTOR KINASE 7 (CARK7) as novel substrates of CPK28. We uncover a role for MAZ and its paralog CARK6 as redundant negative regulators of the immune-triggered oxidative burst. Although they possess all hallmarks of active kinases, we find that MAZ and CARK7 do not possess detectable kinase activity *in vitro,* and that catalytic activity of MAZ is not required for its biological activity *in vivo*. Our work therefore highlights the importance of verifying catalytic activity prior to drawing conclusions about protein function and provides evidence for noncatalytic functions of protein kinases in signaling pathways.

## Results & Discussion

### CPK28 associates with MAZ and CARK7 *in planta*

We recently identified five RLCKs as putative binding proteins of CPK28 in a proteomics screen following affinity purification of CPK28-YFP from *cpk28-1/35S:CPK28-YFP* transgenic lines (Dias *et al*., 2023). This included two members of RLCK subgroup VIII: MAZ and CARK7 (**Figure 1A-B**). Importantly, we did not identify MAZ or CARK7 peptides when we similarly affinity-purified two unrelated plasma membrane resident proteins – Lti6B-GFP (Cutler *et al*., 2000) from Col-0/*35S:Lti6B-GFP* and NSL1-YFP (Holmes *et al*., 2021) from *nsl1-1/35S:NSL1-YFP* (**Figure 1A**) (Dias *et al*., 2023), suggesting specificity. Subgroup VIII RLCKs have been studied in multiple species, including Arabidopsis, soybean, rice, and corn, with demonstrated roles in RLK-mediated signaling networks involved in immunity and development (Liang and Zhou, 2018). For example, the tomato subgroup VIII RLCKs Pto-interacting protein 1 (SlPti1) and SlPti1b are required for the induction of immune-induced oxidative burst and resistance to infection with *P. syringae* pv. *tomato* (Schwizer *et al*., 2017). Moreover, SlPti1 associates with and is phosphorylated by SlPto (Zhou *et al*., 1995). Although originally named PTI1-like kinases in Arabidopsis (PTI1-1 – PTI1-11) (Anthony *et al*., 2006; Huangfu *et al*., 2021), this subfamily was recently renamed CARKs (Zhang *et al*., 2018), and individual members of the subfamily have additional names. The eleven Arabidopsis subgroup VIII RLCKs cluster into three groups (**Figure 1B**) (Herrmann *et al*., 2006). The subgroup VIII-I protein MARIS/CARK4 regulates cell wall integrity sensing resulting in pollen tube and root hair defects (Boisson-Dernier *et al*., 2015; Liao *et al*., 2016). The subgroup VIII-II RLCK CARK1 regulates abscisic acid (ABA) signaling through direct regulation of multiple ABA receptors (Zhang *et al*., 2018; Li *et al*., 2019, 2022). The subgroup VIII-III RLCK CARK8, and to a lesser extent CARK3, CARK7, and MAZ, may be involved in reactive oxygen stress signaling mediated by the kinase OXIDATIVE SIGNAL INDUCIBLE 1 (OXI1) (Anthony *et al*., 2006; Forzani *et al*., 2011). In our previous work, we uncovered that the subgroup VIII-III protein MAZ is a component of stomatal and root development signaling mediated by CLAVATA1 (CLV1) family receptors that perceive CLV3/EMBRYO SURROUNDING REGION-RELATED (CLE) peptides (Blümke *et al*., 2021). Given the broad roles for subgroup VIII RLCKs in cellular signaling, we were motivated to study MAZ, CARK7, and related proteins in more detail.

**Figure 1.**
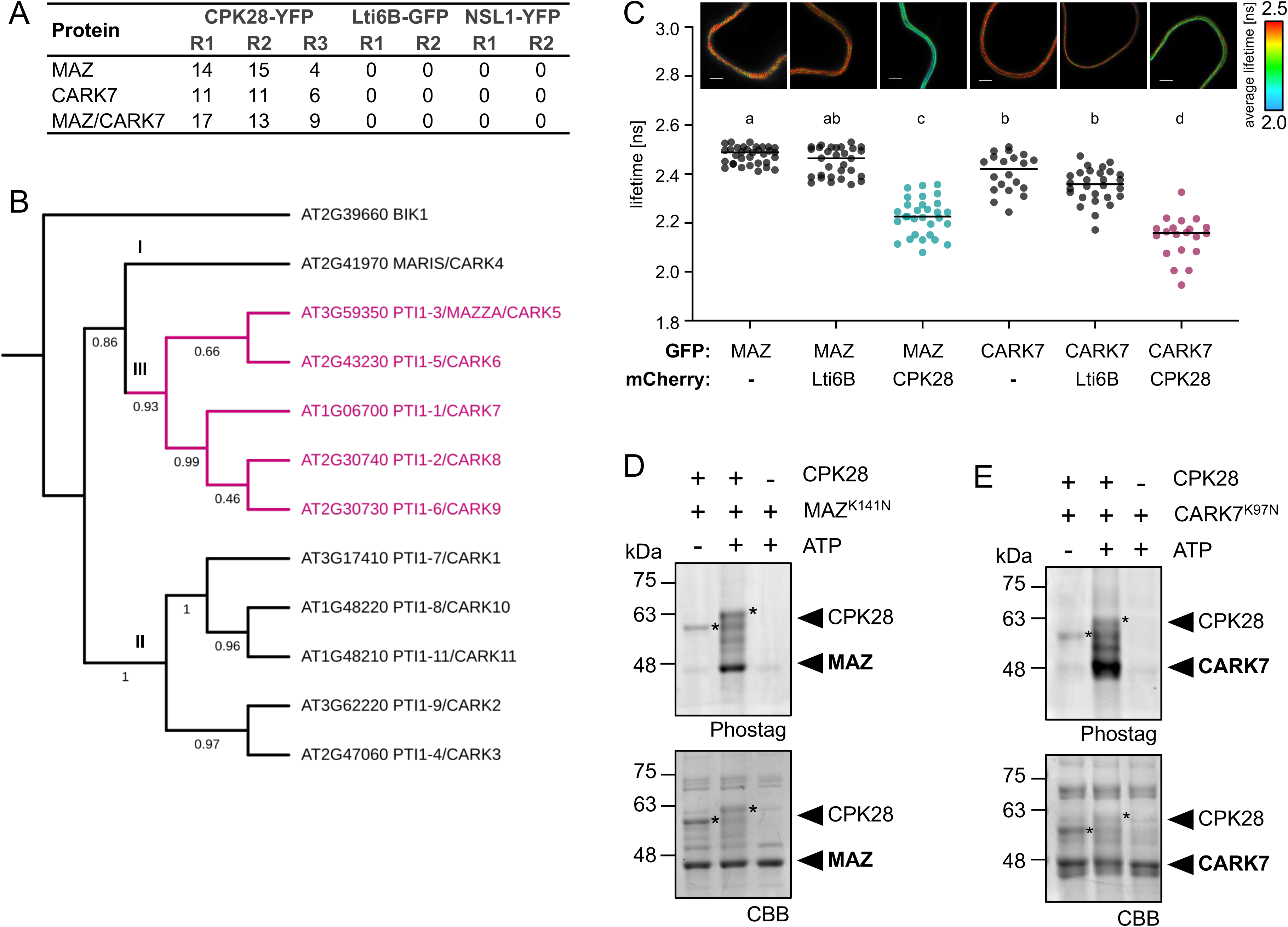
CPK28 associates with and phosphorylates MAZ and CARK7. **(A)** Number of unique peptides matching MAZ and CARK7, as well as peptides that match both MAZ and CARK7, identified by LC-MS/MS after affinity purification of CPK28-YFP, compared to Lti6B-GFP and NSL1-YFP as controls. R1, R2, and R3 refer to independent replicates. The full dataset has been previously published (Dias *et al*., 2023). **(B)** Phylogenetic tree of the Arabidopsis subgroup VIII RLCK family, with BIK1 as an outgroup. The tree was constructed using maximum likelihood with 1000 bootstraps in MEGAX; generated by JM. **(C)** FLIM-FRET analysis of proteins expressed in *N. benthamiana*. The fluorescence lifetime (ns) of MAZ-GFP and CARK7-GFP is reduced only in the presence of CPK28-mCherry and not when expressed alone or in the presence of Lti6B-mCherry. Representative confocal micrographs are displayed above the plot. At least 20 independent cells were measured for each; statistically significant groups are indicated by lower-case letters based on a one-way ANOVA and Tukey’s post-hoc test (*p*<0.002). Data collected by JEM. **(D-E)** *In vitro* kinase assays using CPK28 as the kinase and MAZ^K141N^ **(D)** or CARK7^K97N^ **(E)** as substrates, visualized using Phostag gel staining. Coomassie Brilliant Blue (CBB) indicates loading. These experiments were repeated more than 4 times by TD over the course of a year and representative blots are shown.

To confirm that CPK28 is able to associate with MAZ and CARK7, we first employed split-luciferase complementation analysis (SLCA) using transient expression in *Nicotiana benthamiana*. Here, the N-terminal canonical domain of firefly luciferase (nLuc) is translationally fused to the C-terminus of one protein of interest, while the C-terminal canonical domain of luciferase (cLuc) is N-terminally fused to another protein of interest. If the two proteins associate, a functional luciferase is formed that can produce light if given the substrate luciferin. For these experiments, we used the RLK FERONIA (FER) as a control because it is similarly located at the plasma membrane. We found that co-expressing CPK28-nLuc and cLuc-MAZ reconstituted luciferase function, whereas co-expressing FER-nLuc and cLuc-MAZ did not (**Supplemental Figure S1A**). Similarly, luciferase was reconstituted when we co-expressed CPK28-nLuc and cLuc-CARK7, but not when we co-expressed FER-nLuc and cLuc-CARK7 (**Supplemental Figure S1B**). We next tested if CPK28 can associate with MAZ and CARK7 using fluorescence lifetime imaging-based Förster resonance energy transfer (FLIM-FRET) measurements. In FRET experiments, two proteins of interest are tagged with distinct fluorescent proteins that have overlapping excitation and emission spectra. The phenomenon occurs as a radiation-free energy transfer from a ‘donor’ fluorophore to an ‘acceptor’ fluorophore if they are in close proximity and appropriate orientation. FLIM measures the decrease in the excited-state lifetime of the donor fluorophore when in the presence of an acceptor and can provide information about protein-protein interactions (Spatola Rossi *et al*., 2022). Here, we transiently co-expressed CPK28-mCherry or Lti6B-mCherry with either MAZ-GFP or CARK7-GFP in *N. benthamiana*. We found that the fluorescence lifetime of both MAZ-GFP and CARK7-GFP was reduced in the presence of CPK28-mCherry but not Lti6B-mCherry (**Figure 1C**), suggesting that CPK28 is sufficiently close to MAZ and CARK7 to allow energy transfer. Moreover, confocal micrographs suggest that this association occurs in the plasma membrane, which is where CPK28 and MAZ localize (Monaghan *et al*., 2014; Blümke *et al*., 2021). We conclude that CPK28 is able to associate with both MAZ and CARK7 *in planta*.

### CPK28 phosphorylates MAZ and CARK7 on multiple residues *in vitro*

Several *in vivo* and *in vitro* phosphorylation sites have been mapped on subgroup VIII RLCKs from various phosphoproteomics screens (**Supplemental Figure S2**), suggesting that they may be regulated by phosphorylation. CPK28 is an active protein kinase (Matschi *et al*., 2013; Monaghan *et al*., 2014) that phosphorylates several target proteins (Wang *et al*., 2018*a*; Ding *et al*., 2022; Dias *et al*., 2023) including the subgroup VII RLCK BIK1 (Monaghan *et al*., 2014), which is closely related to MAZ and CARK7. Because they associate, we reasoned that CPK28, MAZ, and CARK7 may engage in trans-phosphorylation. We first tested if CPK28 can phosphorylate MAZ and CARK7. As MAZ and CARK7 are protein kinases, we generated catalytically inactive variants His_6_-MAZ^K141N^ and His_6_-CARK7^K97N^ (in which the ATP-binding Lys has been mutated to Asn) to use as substrates for CPK28. We found that CPK28 was able to phosphorylate both MAZ^K141N^ and CARK7^K97N^ in an ATP-dependent manner, as visualized by Phostag staining (**Figure 1D-E**). To test if MAZ and CARK7 are able to phosphorylate CPK28, we used the catalytically inactive His_6_-CPK28^D188A^ variant (in which a critical Asp in the activation loop has been mutated to Ala) (Matschi *et al*., 2013; Monaghan *et al*., 2014) as substrate. We found that neither MAZ nor CARK7 were able to reciprocally phosphorylate CPK28^D188A^ (**Supplemental Figure S1C-D**). To identify which residues are phosphorylated by CPK28, we performed additional kinase assays using MAZ^K141N^ and CARK7^K97N^ as substrates and analyzed phosphopeptides by liquid chromatography followed by tandem mass spectrometry (LC-MS/MS). We identified nine phosphosites on MAZ and seven phosphosites on CARK7, four of which were in conserved positions on MAZ and CARK7 (**Figure 2A-B**). Interestingly, the phosphosites were located in regions that are considered critical for kinase activation, including in the glycine-rich loop, active site, and in the activation loop (**Figure 2B**). We conclude that CPK28 is able to unidirectionally phosphorylate MAZ and CARK7 on multiple residues.

**Figure 2.**
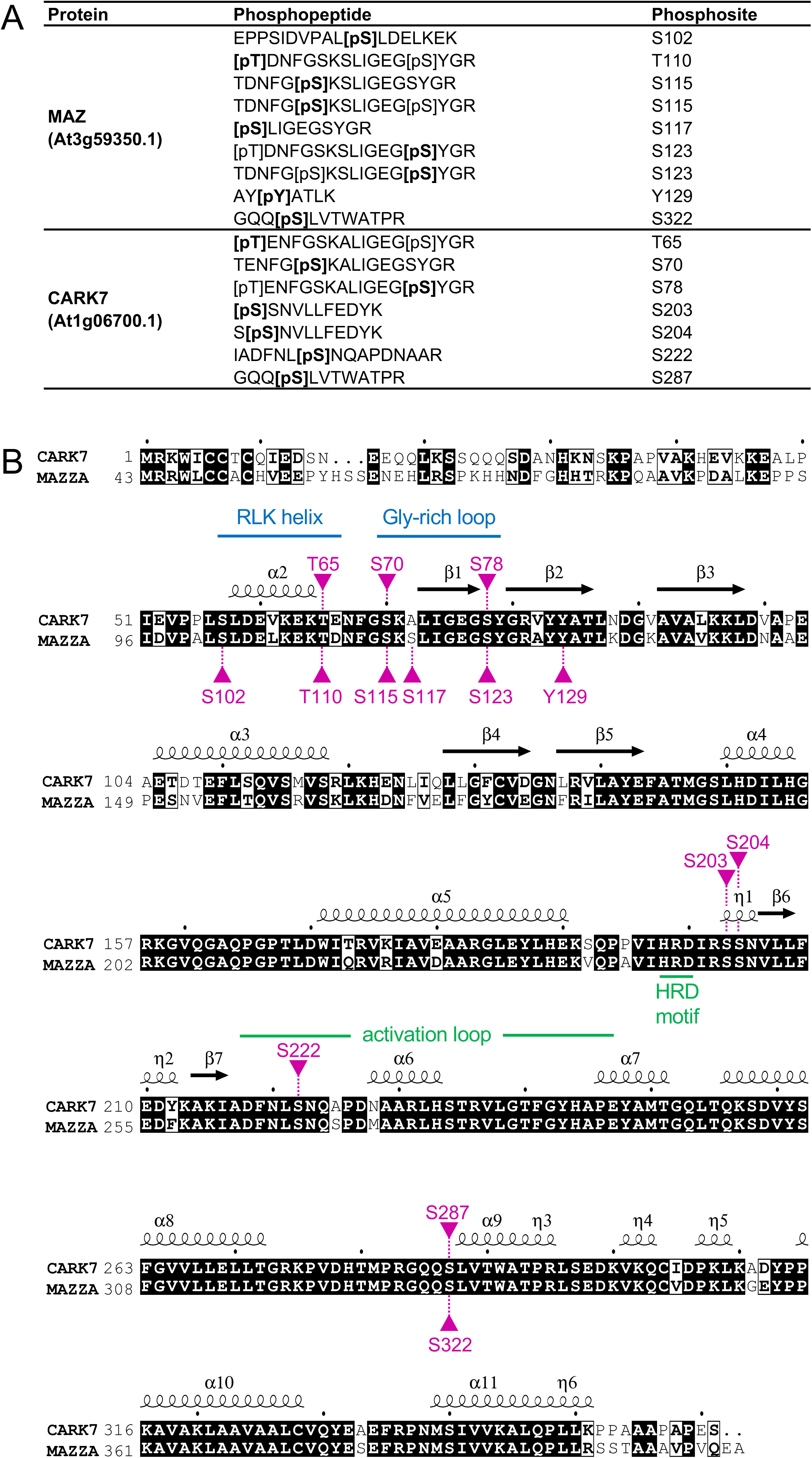
CPK28 phosphorylates MAZ and CARK7 on multiple residues. **(A)** Unique MAZ and CARK7 phosphopeptides following kinase assays with CPK28. Each peptide was present in at least 2/3 independent replicates and not found in control samples with ATP but without CPK28. The bolded sites are the phosphosites and their positions are indicated on the right. Kinase assays performed by TD; trypsin digests and LC-MS/MS performed by MCRG. **(B)** Sequence alignment between MAZ and CARK7 with protein secondary structures indicated as visualized by ESPript (Robert and Gouet, 2014). Residues phosphorylated by CPK28 are indicated in pink (CARK7 sites on top and MAZ sites on the bottom). Analysis by JM.

### MAZ and CARK6 inhibit immune-triggered ROS

Given the documented roles of RLCKs in immune signal transduction (Liang and Zhou, 2018), we sought to assess if subgroup VIII RLCKs play a role in immune signaling. We obtained two insertional mutants in each of *PTI1-3/CARK5/MAZZA* and its paralog *PTI1-5/CARK6,* as well as *PTI1-1/CARK7* and its paralogs *PTI1-2/CARK8* and *PTI1-6/CARK9* (**Figure 3A)**. While we were unable to identify any lines carrying the annotated T-DNA insertion in *cark8-1* (WDL_Hs008-06A)*, cark8-2* (Sail_147C05), or *cark9-2* (Salk_079894), we were able to identify homozygous alleles in all other lines and confirmed that their target genes were downregulated (**Figure 3B-E**). Expecting some genetic redundancy between paralogs, we also generated *maz-1 cark6-2* and *cark7-1 cark9-1* double mutants. Leaf morphology for all single and double mutant lines was indistinguishable from Col-0 wild-type (**Figure 3F**).

**Figure 3.**
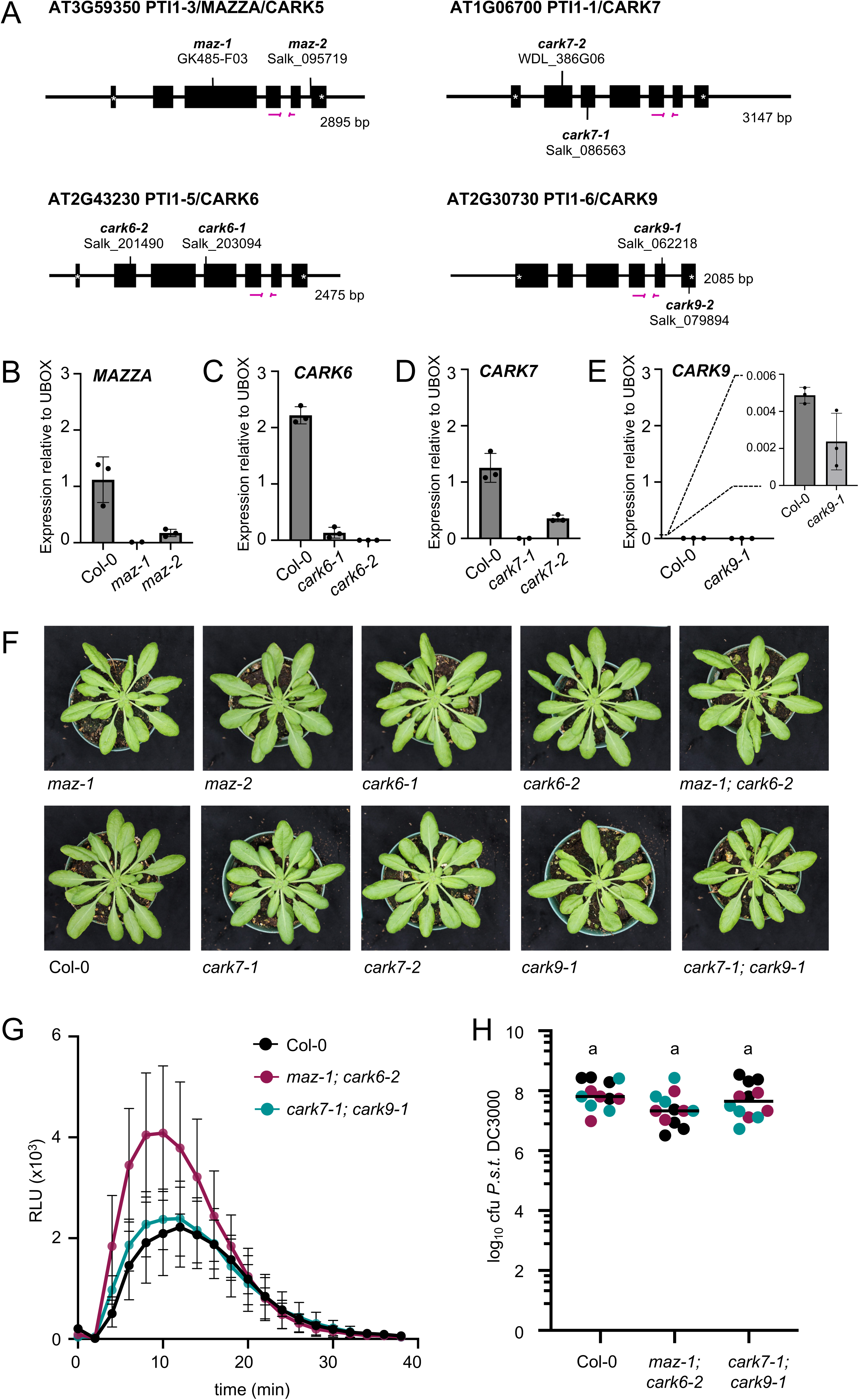
MAZ and CARK6 inhibit immune-triggered ROS. **(A)** Schematic representation, drawn to scale, of Arabidopsis subfamily VIII RLCK genes, indicating exons (boxes), untranslated regions (lines), and the location of T-DNA insertion alleles. Genomic information was retrieved from The Arabidopsis Information Resource (TAIR) by CGF and JM. Lines were genotyped to homozygosity by CGF as outlined in **Supplemental Table S1**. **(B-E)** Quantitative real-time qRT-PCR of MAZ **(B)**, CARK6 **(C)**, CARK7 **(D)**, and CARK9 **(E)**, relative to UBOX. Means for 3 independent biological replicates are shown +/-standard error of the mean. All data collected by CGF. Primers for genotyping and qRT-PCR are provided in **Supplemental Table S1**. **(F)** Photographs of representative plants of each genotype after 5 weeks of growth on soil under short-day conditions. Photographs taken by MGD. **(G)** ROS production measured in relative light units (RLUs) after treatment with 100 nM flg22. Values represent means +/-standard error (n=6 leaf discs). Assays were repeated several times by MGD and JM over multiple years; representative data is shown. **(H)** Growth of *Pseudomonas syringae* pv. *tomato* (*P.s.t.*) isolate DC3000 3 days after syringe-inoculation. Data from 3 independent biological replicates are plotted together, denoted by black, teal, and magenta dots. Values are colony forming units (cfu) per leaf area (cm^2^) from 4-5 samples per genotype (each sample contains 3 leaf discs from 3 different infected plants). The line represents the mean (n=12). Lower case letters indicate that all genotypes are in the same statistical group, as determined by a one-way ANOVA followed by Tukey’s post-hoc test. Data collected by MGD.

A burest of reactive oxygen species (ROS) is readily observable within the first 5-30 min following exposure to immunogenic molecules such as bacterial flagellin (Gómez-Gómez *et al*., 1999). We did not observe any differences in flg22-induced ROS in the *maz-1, maz-2, cark6-1, cark6-2, cark7-1, cark7-2,* or *cark9-1* single mutants, nor in the *cark7-1 cark9-1* double mutant, compared to Col-0 (**Figure 3G; Supplemental Figure S3A-E**). We did, however, observe enhanced flg22-induced ROS in the *maz-1 cark6-2* double mutant (**Figure 3G; Supplemental Figure S3E**). The rapid phosphorylation and activation of MAPKs occurs in parallel with the oxidative burst, also observable within the first 5-30 minutes of exposure to immunogenic peptides (Asai *et al*., 2002; Son *et al*., 2011). We found that the activation of MPK3/6 and MPK4 was similar in Col-0, *maz-1 cark6-2*, and *cark7-1 cark9-1* double mutants **(Supplemental Figure S3F)**, suggesting that subgroup VIII RLCKs do not function in the parallel MAPK pathway. As both ROS and MAPK activation are considered ‘early’ immune responses (Yu *et al*., 2017), we extended our study to also include ‘later’ responses, including both immune-triggered inhibition of seedling growth and resistance to a bacterial pathogen. In Col-0 seedlings, continual exposure to 100 nM flg22 for 10-14 days results in 60-90% reduction in seedling weight (**Supplemental Figure S3G**) (Gómez-Gómez *et al*., 1999). This effect was similar in both *maz-1 cark6-2* and *cark7-1 cark9-1* double mutants (**Supplemental Figure S3G**). In addition, we did not observe any differences between Col-0 and the double mutants when we infected adult plants with the virulent bacterial pathogen *Pseudomonas syringae* pv. *tomato* DC3000 (**Figure 3H**). Together, these data suggest that MAZ and CARK6 inhibit the immune-triggered oxidative burst but that this does not correlate to enhanced resistance against *P.s.t.* DC3000.

Interestingly, while the subgroup VIII RLCKs SlPti1a and SlPTi1b from tomato (Zhou *et al*., 1995; Schwizer *et al*., 2017; Giska and Martin, 2019), NbPti1b and NbPti1c from *N. benthamiana* (Schwizer *et al*., 2017), and CsPti1-L from cucumber (Oh *et al*., 2014) promote immune signaling, the subgroup VIII RLCK OsPti1a negatively regulates immunity in rice (Takahashi *et al*., 2007; Matsui *et al*., 2010). Overexpression of OsPti1a reduces resistance to rice blast fungus, while OsPti1a knockouts are characterized by chlorotic lesions and other hallmarks of deregulated immune signaling (Takahashi *et al*., 2007). The hyperimmunity and enhanced resistance observed in *Ospti1a* knockouts can be suppressed by silencing *OsRAR1* (*REQUIRED FOR AVRB-DEPENDENT SUSCEPTIBILITY AND CHLOROSIS*), a well-known downstream component of NLR signaling (Shirasu, 2009). Therefore it is also possible that OsPti1a is guarded by an NLR (Takahashi *et al*., 2007; Schwizer *et al*., 2017) that is present in rice but absent in solanaceous plants (Schwizer *et al*., 2017). Our finding that MAZ and CARK6 negatively regulate immune triggered ROS suggests roles for subgroup VIII as negative regulators of defense. Although we were unable to obtain *cark8* mutants, previous work has shown that CARK8 activity increases in response to stress signals including phosphatidic acid, hydrogen peroxide, and the bacterial immunogenetic elicitors xylanase and flagellin (Anthony *et al*., 2006), which may suggest a role in immune signaling. As this subfamily is large in Arabidopsis, and as genetic redundancy is well-known amongst other RLCKs (Rao *et al*., 2018), it is likely that a quintuple mutant lacking MAZ, CARK6, CARK7, CARK8, and CARK9 may reveal stronger immune phenotypes. We propose this as an avenue for future analysis of this gene family.

### MAZ and CARK7 are catalytically inactive *in vitro*

To assess if MAZ and CARK7 possess catalytic autophosphorylation activity, we performed *in vitro* kinase assays using recombinant His_6_-MAZ and His_6_-CARK7. We included either GST-BIK1 or His_6_-CPK28 as technical controls for our assays. Although we could readily detect autophosphorylation on BIK1, we were unable to detect autophosphorylation on MAZ or CARK7 when we supplied the kinases with divalent magnesium or manganese as cofactors along with [γ32P]-ATP (**Figure 4A**). Because some protein kinases are inhibited by phosphorylation, we next expressed and purified the recombinant proteins from an *E. coli* strain that expresses λ-phosphatase. However, we could again not detect any *in vitro* autophosphorylation for both MAZ or CARK7 (**Figure 4B**). Some protein kinases do not autophosphorylate but can transphosphorylate substrates. We therefore assessed if MAZ or CARK7 are able to trans-phosphorylate Histone 3S (a highly phosphorylatable protein considered a universal substrate for protein kinases), however we once again could not detect any activity (**Figure 4C**). We note that a previous study reported that both MAZ and CARK7 (as well as CARK2, CARK4, and CARK11) are able to phosphorylate multiple ABA receptors (Li *et al*., 2022), however in our view these assays were not adequately controlled and are hard to interpret. The CARKs and ABA receptors were combined in the presence or absence of ATP and phosphorylation was visualized using an anti-pThr antibody. Immunoreactive bands are clearly visible for the ABA receptors only in the presence of both CARKs and ATP, however there is no control with the receptors in the presence of ATP and absence of CARKs. We find it difficult to reconcile differences between our studies. We also note that another study showed that CARK7, CARK8, and MAZ exhibit low levels of autophosphorylation (Anthony *et al*., 2006). However, those experiments were performed following transfection of Arabidopsis protoplasts, which may not faithfully reflect autophosphorylation. In the same study, the authors showed that CARK7 and CARK8 can be phosphorylated by OXI1 (Anthony *et al*., 2006), a protein kinase involved in signaling events broadly mediated by reactive oxygen species (Rentel *et al*., 2004).

**Figure 4.**
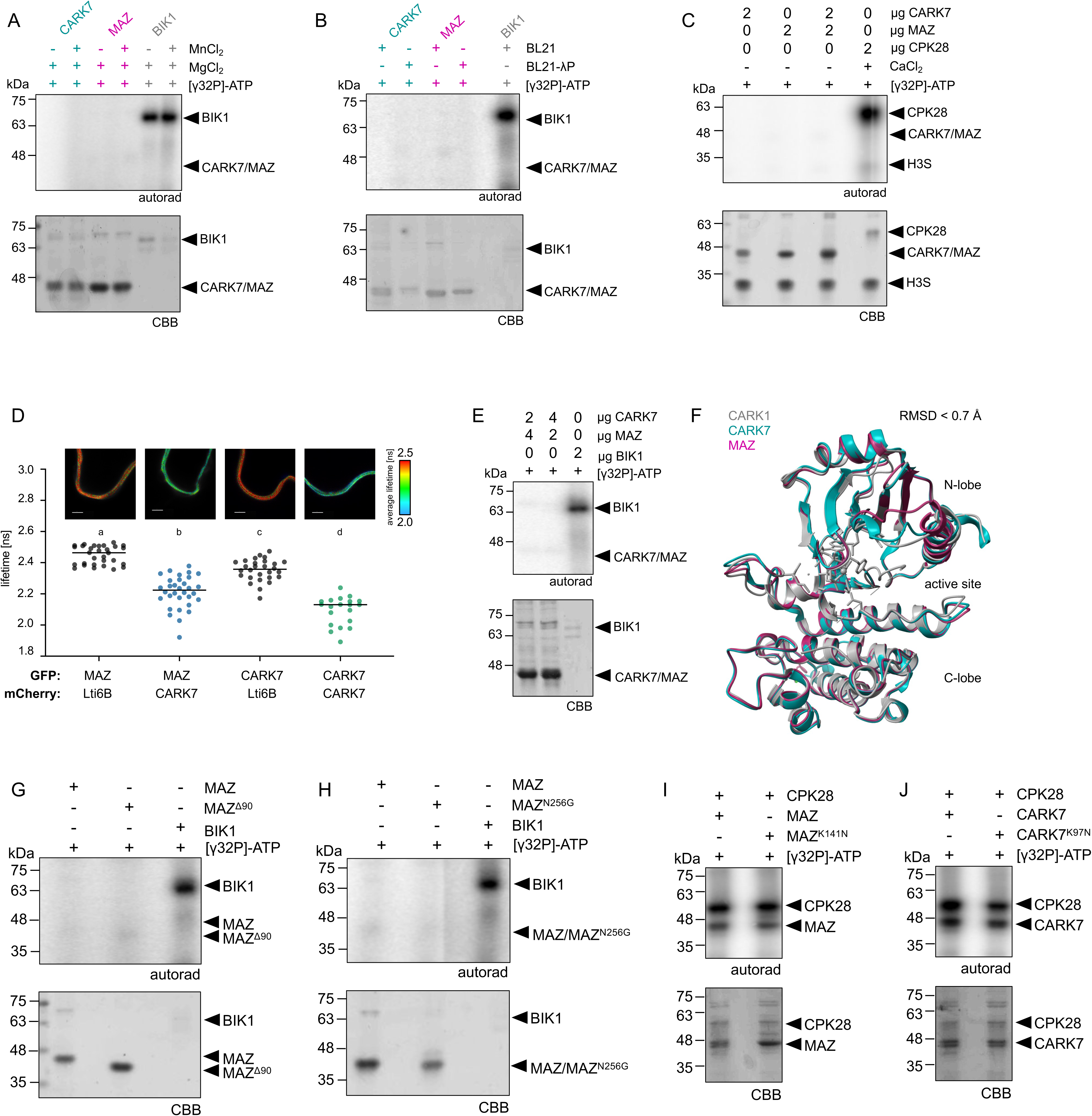
MAZ and CARK7 are catalytically inactive. **(A-B)** *In vitro* autophosphorylation assays using His_6_-MAZ and His_6_-CARK7 alongside GST-BIK1 as a control. Experiments in **(A)** compare the utility of Mn^2+^ or Mg^2+^ as co-factors, while experiments in **(B)** compare proteins purified from *E. coli* BL21 (may already be phosphorylated) with those purified from an *E.coli* BL21 strain expressing λ-phosphatase (to dephosphorylate proteins). Autoradiographs (autorad) indicate incorporation of γP32 and protein loading is indicated by post-staining the membranes with Coomassie Brilliant Blue (CBB). These assays were repeated more than three times by TD; representative experiments are shown. **(C)** *In vitro* transphosphorylation assays between His_6_-MAZ and His_6_-CARK7 and the universal substrate H3S, alongside transphosphorylation of H3S by His_6_-CPK28. Autorad indicates incorporation of γP32 and protein loading is indicated with CBB. These assays were repeated more than three times by TD; representative experiments are shown. **(D)** FLIM-FRET analysis of proteins expressed in *N. benthamiana*. The fluorescence lifetime (ns) of MAZ-GFP and CARK7-GFP is reduced only in the presence of CARK7-mCherry and not when expressed in the presence of Lti6B-mCherry. Representative confocal micrographs are displayed above the plot. At least 20 independent cells were measured for each; statistically significant groups are indicated by lower-case letters based on a one-way ANOVA and Tukey’s post-hoc test (p<0.0001). Data collected by JEM. **(E)** *In vitro* transphosphorylation assays between His_6_-MAZ and His_6_-CARK7. Autorad indicates incorporation of γP32 and protein loading is indicated with CBB. These assays were repeated more than three times by TD; representative experiments are shown. **(F)** Protein structure overlays comparing the AlphaFold-predicted structures of CARK7 and MAZ with the solved structure of CARK1 (PBD: 5XD6), determined using the matchmaker command in ChimeraX (Pettersen *et al*., 2021). The root-mean-square deviation (RMSD) of <0.7 Å indicates a high level of similarity. **(G-H)** *In vitro* autophosphorylation assays using His_6_-MAZ^Δ90^ **(G)** and His_6_-MAZ^N256G^ **(H)** alongside GST-BIK1 as a control. Autorad indicates incorporation of γP32 and protein loading is indicated with CBB. Data in **(G)** was collected by TD using protein purified by VNM, while data in **(H)** was collected by TD using protein purified by TD. Assays were repeated more than three times and representative experiments are shown. **(I-J)** *In vitro* trans-/auto-phosphorylation assays between His_6_-CPK28 and His_6_-MAZ or His_6_-MAZ^K141N^ **(I)**, or His_6_-CARK7 or His_6_-CARK7^K97N^ **(J).** Autorad indicates incorporation of γP32 and protein loading is indicated with CBB. These assays were repeated more than three times by TD; representative experiments are shown.

Previous work has shown that many CARKs are able to associate with themselves and each other (Li *et al*., 2022), suggesting that homo– and/or hetero-dimerization may be an important feature of subgroup VIII RLCKs. For example, MAZ can associate with itself (Blümke *et al*., 2021) as well as CARK6 (Li *et al*., 2022), and CARK7 can associate with CARK11 (Li *et al*., 2022). To test if MAZ can associate with CARK7, we again employed SLCA and found that co-expression of CARK7-nLuc and cLuc-MAZ successfully reconstituted luciferase function, whereas co-expression of FER-nLuc and cLuc-MAZ did not (**Supplemental Figure S4A**). Similarly, in FLIM-FRET experiments, we found that the lifetime of MAZ-GFP was reduced when co-expressed with CARK7-mCherry, but not when co-expressed with Lti6B-mCherry (**Figure 4D**). In addition, we observed CARK7 self-association in both SLCA (**Supplemental Figure S4B**) and FLIM-FRET (**Figure 4D**) experiments. We conclude that CARK7 and MAZ are able to associate with themselves and each other, possibly forming homo– and hetero-dimers, which may reflect a general feature of this protein family. To test the hypothesis that heterodimer formation may regulate MAZ and CARK7 catalytic activity, we next conducted kinase assays with both proteins together. However, we were again unable to detect phosphoryl transfer (**Figure 4E**). Overall, we conclude that MAZ and CARK7 are catalytically inactive *in vitro*.

These results were unexpected since most studied RLCKs possess kinase activity (Liang and Zhou, 2018). We therefore interrogated the sequences of subgroup VIII RLCKs in detail in an attempt to uncover why they are unable to catalyze phosphorylation *in vitro.* A survey of the literature revealed that while several subgroup VIII RLCKs from multiple species do autophosphorylate *in vitro* (Zhou *et al*., 1995; Anthony *et al*., 2006; Herrmann *et al*., 2006; Zou *et al*., 2006; Takahashi *et al*., 2007; Forzani *et al*., 2011; Oh *et al*., 2014; Zhang *et al*., 2018; Giska and Martin, 2019; Li *et al*., 2019; Peng *et al*., 2022), others do not (Staswick, 2000; Tian *et al*., 2004; Liao *et al*., 2016). We therefore decided to take a comparative approach. The protein structures predicted by AlphaFold (Jumper *et al*., 2021) for MAZ and CARK7 indicate that both are likely to fold into canonical protein kinases with well-structured N– and C-lobes. We superimposed the predicted structures for MAZ and CARK7 with the experimentally-determined structure for CARK1 (PBD: 5XD6) which has autophosphorylation activity (Zhang *et al*., 2018), and found a high level of similarity, with minimal root mean square deviations less than 0.7 Å (**Figure 4F**). The only differences we observed were in intrinsically disordered or low-order regions such as the activation loops which have low confidence prediction values and were missing in the CARK1 structure (Zhang *et al*., 2018). Based on this comparison, we surmise that MAZ and CARK7 are likely to fold into proteins that are structurally similar to CARK1. In this regard, it is interesting to note that homo– and hetero-dimerization of CARK1 and CARK3, as well as the association between CARK1 and ABA receptors, is dependent on the presence of catalytic residues (Zhang *et al*., 2018; Li *et al*., 2019, 2022). This may suggest that at least some subgroup VIII RLCKs adopt protein conformations that are sensitive to changes in the catalytic cleft.

Like many RLCKs, MAZ and CARK7 contain a central protein kinase domain flanked by variable N– and C-terminal domains. In the specific case of MAZ and CARK7, the C-terminal domain is very short, but the N-terminal domains are longer (∼100 in MAZ (Blümke *et al*., 2021) and ∼50 aa in CARK7) and are predicted to be intrinsically disordered. We hypothesized that the N-terminal extensions may inhibit kinase activity in some way. Using MAZ as a case study, we generated a mutant variant translationally fused to His_6_ but lacking the majority of the N-terminal domain (starting at L90 following M1; 10 amino acids N-terminal to the canonical kinase domain). We were again unable to detect any autophosphorylation on MAZ^Δ90^ (**Figure 4G**), arguing against an N-terminal auto-inhibition mechanism.

We next generated a multiple sequence alignment of the entire subgroup VIII RLCK subfamily from plants spanning the plant kingdom: dicotyledonous angiosperms *Amborella trichopoda* (3 members), *Arabidopsis thaliana* (11 members), *Solanum lycopersicum* (7 members), *Glycine max* (11 members), and *Cucumis sativus* (3 members); monocotyledonous angiosperms *Oryza sativa* (10 members) and *Zea mays* (13 members); moss *Physcomitrium patens* (2 members), and liverwort *Marchantia polymorpha* (1 member). In all cases, we could identify key features of active protein kinases (Nolen *et al*., 2004): in the N-lobe, we identified the glycine-rich loop and the ATP-binding lysine; in the C-lobe, we identified the ‘HRD’ motif and the critical Asp residue of the ‘DFG’ motif in the activation loop. We noted, however, that all subgroup VIII RLCKs do not possess typical DFG motifs, as the Gly is either replaced by Asn or Asp, resulting in non-canonical DFN or DFD motifs that correspond to phylogenetic clusters within the subgroup VIII RLCK family (**Supplemental Figure S5**). Subgroup VIII-III RLCKs, including MAZ and CARK7, contain DFN. For many protein kinases, the precise positioning of the DFG-Phe is critical for enzyme activation as it subsequently allows for the correct positioning of the DFG-Asp to bind ATP in the active site (Ung and Schlessinger, 2015). Both Asn and Asp are larger amino acids than Gly, which may result in steric hindrance of the neighbouring Phe in the DFN and DFD variants. We therefore generated a variant of MAZ with a canonical DFG motif, His_6_-MAZ^N256G^, and assessed autophosphorylation *in vitro.* However, we were once again unable to detect any activity (**Figure 4H**).

Because CPK28 phosphorylates MAZ and CARK7 on residues known to be critical for kinase activation (**Figure 2**), we hypothesized that they may require transphosphorylation for activation. We therefore performed additional kinase assays using MAZ or CARK7 as substrates for CPK28 in comparison to MAZ^K141N^ or CARK7^K97N^. We reasoned that if MAZ or CARK7 required phosphorylation by CPK28 for activation, we should observe an increase in the incorporation of [γ32P]-ATP on the wildtype variants compared to the catalytically inactive variants, reflecting both transphosphorylation events by CPK28 as well as autophosphorylation events by MAZ or CARK7. However, we did not observe any noticeable difference between the substrates (**Figure 4I-J**). This suggests that phosphorylation by CPK28 is not sufficient to activate the enzymatic function of MAZ or CARK7. Overall, after testing several hypotheses, we conclude that neither MAZ nor CARK7 display catalytic activity *in vitro*.

### The catalytic activity of MAZ is not required for its role in the CLV3p signaling pathway

Although we could not detect any activity *in vitro,* it is possible that MAZ, CARK7, and other subgroup VIII RLCKs are active kinases *in vivo* under specific contexts. For example, they may require cellular cues or activation by upstream protein kinases or binding partners similar to mitogen-activated or cyclin-dependent protein kinases (MAPKs and CDKs). If this is the case, it is reasonable to expect their catalytic activity to be essential to carry out their biological roles. Although our analysis above indicates redundant roles between MAZ and CARK6 in immune-triggered ROS (**Figure 3G; Supplemental Figure S3E**), we recently showed that *maz-1* single mutants are partially resistant to CLV3p-triggered root meristem differentiation, which is genetically complemented in both *maz-1/pUBQ10:MAZ-GFP* and *maz-1/pMAZ:MAZ-mCherry* transgenic lines (Blümke *et al*., 2021). To test if kinase activity is required for the biological function of MAZ, we generated two independent and homozygous *maz-1/p35S:MAZ^K141N^-GFP* transgenic lines. Both lines grew similarly to Col-0 with no aberrant leaf morphology or any obvious growth defects (**Figure 5A**), and we verified that the transgenic lines express MAZ^K141N^-GFP at the expected size of ∼74 kDa **(Figure 5B)**. Interestingly, in both cases, expression of *p35S:MAZ^K141N^-GFP* was able to fully restore sensitivity to CLV3p in *maz-1* in a manner comparable to *pUBQ:MAZ-GFP* and *pMAZ:MAZ-mCherry* (**Figure 5C**). These results suggest that the catalytic activity of MAZ is dispensable for its role in CLV1f-mediated signaling. As we were unable to assess if catalytic activity of MAZ is required for its role in immune-triggered ROS using these lines, it is unclear if MAZ and/or CARK6 require catalytic activity in immune signaling. Nevertheless, these results demonstrate that the biological function of MAZ involves a noncatalytic mechanism.

**Figure 5.**
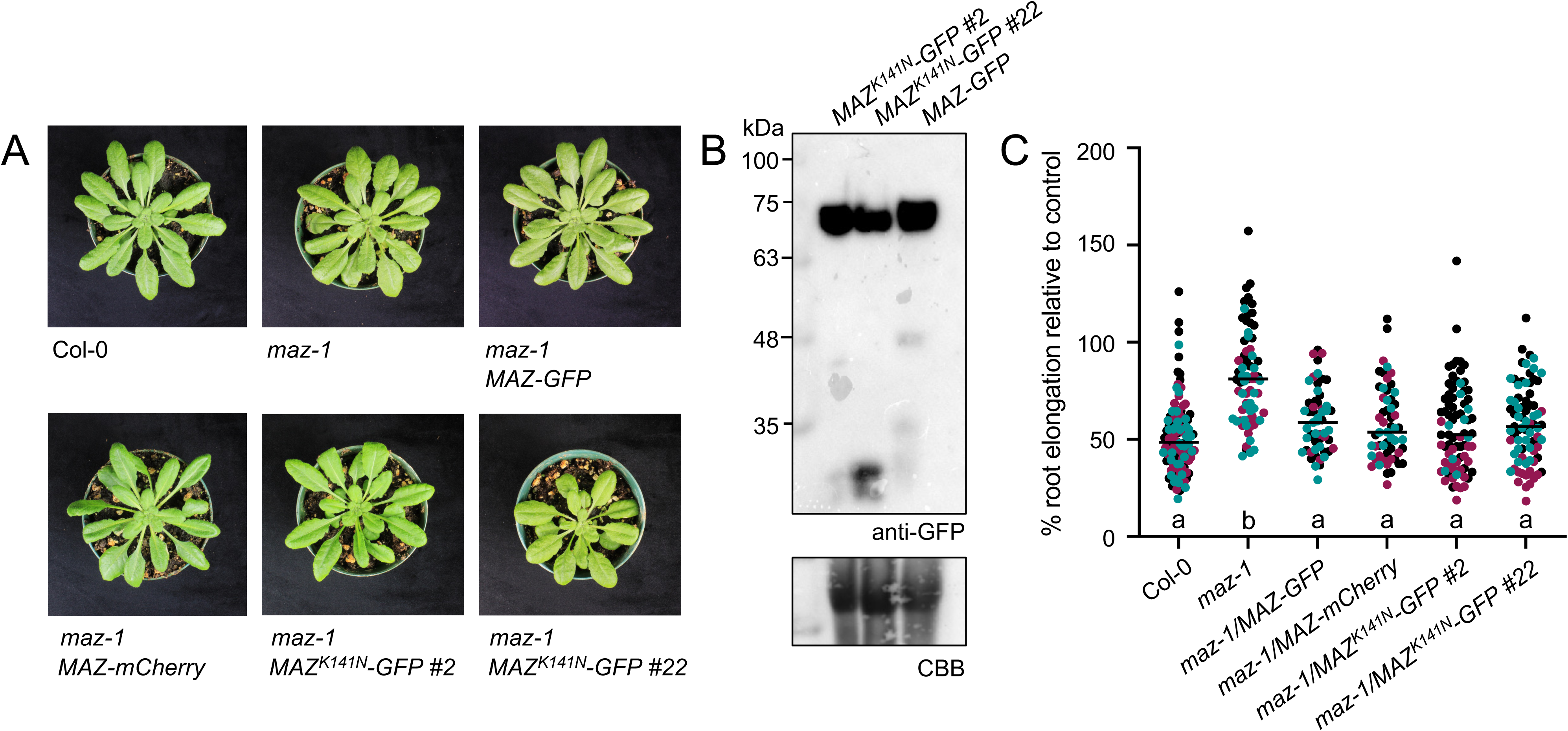
The catalytic activity of MAZ is not required for its role in the CLV3p signaling pathway. **(A)** Photographs of representative plants of each genotype after 5 weeks of growth on soil under short-day conditions. Transgenic lines were generated and photographed by MGD. **(B)** MAZ^K141N^-GFP (∼74 kDa) migrated to its expected size in the *maz-1/35S:MAZ^K141N^-GFP* transgenic lines. Coomassie Brilliant Blue (CBB) of RuBisCO indicates loading. This experiment was repeated twice by MGD with identical results. **(C)** Root length of 55-119 10-day-old seedlings grown on 0.5x MS agar plates supplemented with 10 nM CLV3p, relative to mean root length of 10-day-old seedlings grown on 0.5x MS agar with no peptide. Data from 3 independent biological replicates are plotted together, denoted by black, teal, and magenta dots. Lower case letters indicate statistically significant groups, determined by a one-way ANOVA followed by Tukey’s post-hoc test (p<0.0001). Data collected by MGD.

Noncatalytic roles for protein kinases are now well recognized, including roles as allosteric regulators and protein scaffolds (Kung and Jura, 2016). Recently, the LRR-RLK EFR (ELONGATION FACTOR TU RECEPTOR) was shown to undergo phosphorylation-mediated conformational changes that allow for the allosteric activation of its co-receptor BAK1 (Bender *et al*., 2021; Mühlenbeck *et al*., 2024). Although EFR is an active kinase *in vitro*, mutant variants that impair catalytic activity *in vivo* remain capable of initiating immune signaling (Bender *et al*., 2021). The current model suggests that phosphorylation within the activation loop stabilizes an active conformation of EFR that allows it to allosterically activate BAK1 to initiate immune signaling (Mühlenbeck *et al*., 2024). Although we do not have any evidence to support a similar model for subgroup VIII RLCKs, it is possible that phosphorylation by upstream kinases such as CPK28 could help stabilize conformations of MAZ and CARK7 that are capable of allosterically regulating downstream partners, possibly also acting as scaffolds to enable or coordinate molecular complex formation.

## Methods

### Germplasm and plant growth conditions

*A. thaliana* and *N. benthamiana* plants were grown in the Queen’s Phytotron Facility or in phytochambers at Heinrich Heine University. For aseptic growth, *A. thaliana* seeds were surface sterilized using a 40% bleach solution and sown onto petri plates with 0.5x Murashige and Skoog (MS) media containing 0.8% agar. Afterward, they were subjected to cold stratification in darkness at 4°C for 3-5 days before being exposed to light. For plants grown in soil, seeds were directly sown onto potting soil (Sungro Sunshine Mix 1 or Fafard’s Agro G6 with Coco). Seedlings were later transplanted either individually into 3” pots or six plants per 8” pot two weeks after sowing. Controlled growth chambers (BioChambers and Conviron) were utilized for plant cultivation, maintaining a 10-hour light and 8-hour dark cycle at 22°C, with 30% relative humidity and a light intensity of 150 µE m^2^ s^-1^. Plants were watered from the top as needed, typically every other day, and fertilized every two weeks with a solution containing 1.5 g/L of 20:20:20 N:P:K. *N. benthamiana* seeds followed a similar procedure, sown on potting soil and transplanted individually, but were cultivated in a dedicated growth chamber (Conviron) with a 16-hour light and 8-hour dark cycle, receiving weekly fertilization. Additionally, mite bags containing *Amblyseius swirskii* (Koppert) were introduced bi-weekly to each plant tray to prevent pest infestations.

Information on all germplasm used in this study is outlined in detail in **Supplemental Table S1**. Segregating T-DNA insertion lines were obtained from the Arabidopsis Biological Resource Centre (ABRC) and genotyped using gene-specific primers in standard polymerase chain reactions (PCR). To determine gene expression, we extracted RNA using Aurum Total RNA Mini Kit (BioRad), according to the manufacturer’s instructions. Reverse transcription (RT) was achieved using Superscript III (Invitrogen), according to the manufacturer’s instructions. Quantitative RT-PCR was performed using Sso Advanced Universal SYBR Green Supermix (BioRad) according to the manufacturer’s directions, on a CFX96 Touch Real-Time PCR Detection System (BioRad), using primers specific to each gene as outlined in **Supplemental Table S1**. Higher-order mutants were generated by crossing and genotyping to homozygosity in the F_2_ and F_3_ generations. Transgenic plants were generated by floral dip with *Agrobacterium tumefaciens* GV3101 carrying a suitable binary vector for selection, as previously described (Clough and Bent, 1998). Independent lines were followed through four generations and genotyped to homozygosity based on 100% resistance to the selection marker in the T_3_ generation. All details relating to the CPK28 proteomics screen are described in full elsewhere (Dias *et al*., 2023).

### Molecular cloning

Detailed information about all vectors used in this study, including previously published vectors, can be found in **Supplemental Table S1.** Recombination of entry vectors into Gateway-compatible binary vectors pK7FWG2 (Karimi *et al*., 2002), pABindGFP (Bleckmann *et al*., 2010), or pABindmCherry (Bleckmann *et al*., 2010) was achieved using Gateway LR Clonase II (Invitrogen) according to the manufacturer’s instructions. Directional cloning by digestion-ligation into pCAMBIA1300-nLuc and pCAMBIA1300-cLuc (Chen *et al*., 2008) was achieved using PCR amplification with Q5 Taq polymerase, construct-specific endonucleases and T4 DNA ligase (all enzymes from NEB BioLabs), according to the manufacturer’s directions. Successful assembly of all plasmids was verified using Sanger sequencing (Centre for Applied Genomics, Toronto ON, Canada) or whole-plasmid sequencing (Plasmidsaurus, Eugene OR, USA). For recombinant protein production in *Escherichia coli*, gene coding regions were synthesized and cloned into pET28a+ (Novagen) by Twist BioSciences (San Francisco, USA).

### Split-luciferase complementation

*A. tumefaciens* GV3101 carrying plasmids for split-luciferase complementation was infiltrated into fully expanded leaves of 3-4 week old *N. benthamiana* plants alongside the viral suppressor P19 (Voinnet *et al*., 2003). Leaf discs were harvested after 3 days and exposed to 1 mM luciferin (Gold Biotechnology) for 10 minutes in the dark, followed by recording light emission (integration time of 1 sec) using the LUM module in a SpectraMax Paradigm Multi Mode Microplate Reader.

### FLIM-FRET

Fluorescence lifetime was measured on a Zeiss LSM 780 confocal microscope (40× water immersion objective, Zeiss C-PlanApo, NA 1.2). For TCSPC, a PicoQuant Hydra Harp 400 (PicoQuant, Berlin, Germany) was used. Photon counting was performed with picosecond resolution. GFP was excited with a 485 nm pulsed polarized laser (LDH-D-C-485, 32 MHz, PicoQuant, Berlin, Germany). mCherry was excited with a 561 nm laser with 1% laser power. Laser power at the objective lens was adjusted to 1 µW for the 485 nm laser. Light, emitted from the sample, was separated by a polarizing beam splitter before photons were selected with a band-pass filter. For GFP, a 520/30 band-pass filter; and for mCherry, a 607/70 band-pass filter was used. When GFP served as a donor, a LP610 beam splitter was used. Photons were detected with Tau-SPADs (PicoQuant, Berlin, Germany). Images were acquired at zoom 8 resolution of 256 x 256 pixels with a pixel size of 0.1 µm and a pixel dwell time of 12.54 µs and laser repetition rate of 32 MHz. Photons were collected over 40 frames. To avoid pileup effects, nuclei containing high donor concentrations were avoided. Before image acquisition, the system was calibrated. For this, the objective was adjusted to reach a maximal count rate. FCS curves of Rhodamine110 dye and water were acquired to monitor the system function. Internal response functions for each laser were determined by measuring the fluorescence decay of quenched erythrosine in saturated KI using the same hardware settings as for the FRET pair.

The fluorescence decays of selected ROIs in the FLIM image were analyzed with the SymPhoTime FLIM analysis software (SymPhoTime 64, version 2.4; PicoQuant, Berlin, Germany). TCSPC bins of channel 1 and 2 (parallel and perpendicular light) were binned by eight resulting in a bin width of 8 ps. Nuclei were selected by hand using the ROI tool. Chloroplasts and pixels above the pile-up limit (10% of the laser repetition rate) were manually removed. Decays from donor only samples were fitted with the FLIM analysis tool (Fitting model: n-exponential reconvolution). Judged by fitting residuals and Chi-square-test, one lifetime (model parameter n=1) was needed to fit donor only decays containing GFP.

FRET samples (containing GFP and mCherry) as well as donor only samples (only GFP) were fitted using the LT FRET image analysis tool. Parameter “τ*_Donor_*” was fixed as the average donor only lifetime measured beforehand in the FLIM analysis. The second lifetime parameter “τ*_FRET_*” corresponds to the FRET fraction of the sample and was fitted within limits corresponding to 10% and 80% of the average lifetime acquired in the donor only samples. The lower limit of the amplitude of the FRET fraction “*A_FRET_*” was set to 0.0001 to avoid fitting of negative amplitudes in donor only samples and samples containing Lti6B-mCherry. Average amplitude weighted lifetimes were calculated as the sum of each lifetime component (τ*_FRET_* and τ*_donor_*) weighted by their respective amplitude:

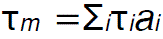

### Immune and growth assays

Immune-triggered ROS, seedling growth inhibition, MAPK activation, and infection with *Pseudomonas syringae* pv. *tomato* DC3000 were performed as described previously (Monaghan *et al*., 2014; Bredow *et al*., 2019; Dias *et al*., 2023). CLV3p-induced root inhibition was performed as described previously (Blümke *et al*., 2021). The 22 amino acid flg22 peptide (Gómez-Gómez *et al*., 1999) was synthesized by EZ Biotech (Indiana USA), and the 13 amino acid CLV3p peptide (Kondo *et al*., 2006) was synthesized by LifeTein (New Jersey USA).

### Recombinant protein purification and *in vitro* kinase assays

All proteins were expressed and purified from *E. coli* strain BL21 or BL21-λP cells as recently described (Dias *et al*., 2023), using the constructs outlined in **Supplemental Table S1**. *In vitro* kinase assays using Phospho-tag gel stain (APB Bio) were performed as per the manufacturer’s instructions and as previously described (Bredow *et al*., 2021). *In vitro* kinase assays using γP32-ATP were performed as recently described (Dias *et al*., 2023), using 2 μg kinase in a buffer containing 50 mM Tris-HCl (pH 8.0), 25 mM MgCl_2_ and/or 25 mM MnCl_2_, 5 mM DTT, 5 μM ATP, and 0.5-2 μCi γP32-ATP. The buffer used in assays with His_6_-CPK28 contained 500 μM CaCl_2_ and no MnCl_2_. Histone 3S from calf thymus was used as a universal substrate in some reactions (Sigma Aldrich H5505). All reactions were incubated for 60 minutes at 30°C. Reactions were stopped by adding 6× Laemmli buffer and heating at 80°C for 5 min. Proteins were separated by 10% SDS-PAGE. The gels were sandwiched between two sheets of transparency film, exposed to a storage phosphor screen (Molecular Dynamics) overnight, and visualized using a Typhoon 8600 Imager (Molecular Dynamics/Amersham). Gels were post-stained with Coomassie Brilliant Blue (CBB) R-250 (MP Biomedicals) or SimplyBlue SafeStain (Invitrogen; CBB G-250) and scanned.

### Phosphoproteomics

*In vitro* kinase assays were performed using 2 μg His_6_-CPK28 as the kinase and 4 μg of either His_6_-MAZ^K141N^ or His_6_-CARK7^K97N^ as substrates in a buffer containing 25 mM Tris-HCl (pH 8.0), 10 mM MgCl_2_, 100 μM CaCl_2_, 1 mM DTT, 1mM PMSF, and 100 μM ATP. Control reactions included His_6_-MAZ^K141N^ or His_6_-CARK7^K97N^ with no kinase. All reactions were incubated for 60 minutes at 30°C and stopped by adding 6× Laemmli buffer and heating at 80°C for 5 min. Protein separation was carried out using 10% Mini-PROTEAN TGX Precast Gels (BioRad 456-1036), followed by post-staining with SimplyBlue SafeStain (Invitrogen; CBB G-250). Bands corresponding to the size of His_6_-MAZ^K141N^ or His_6_-CARK7^K97N^ were excised using sterile razors, and washed with ethanol and sterile ddH_2_0.

SDS-PA gel slices containing immobilized proteins were destained four times with 50% acetonitrile (ACN) in 100 mM tetraethylammonium bromide (TEAB) at 37°C for 10 min. Destain solution was removed and gel pieces were washed with 100 mM TEAB at 37°C for 10 min. Gel pieces were dehydrated by incubating in 100% ACN at room temperature (RT) for 10 min. Gel pieces were then fully dried at 37°C for 5 minutes. Cysteine residues were reduced by 10 mM dithiothreitol solution in 100 mM TEAB for 45 min at 37°C and then alkylated with 55 mM iodoacetamide in 100 mM TEAB buffer in the dark for 1 h at room temperature. Gel pieces were then washed in 50 mM TEAB buffer for 10 min, dehydrated by incubating twice in 100% ACN at RT and then fully dried at 37°C for 5 min. Gel pieces were rehydrated by adding 6 ng/μL trypsin (Promega SequencingGrade – V5113) in 50 mM TEAB. Peptides were digested for 16 h at 37°C shaking at 150 rpm. In-solution tryptic peptides were retained in a separate tube, and in-gel digested peptides were further extracted by adding 1% formic acid, 2% acetonitrile in 100 mM TEAB and allowed to incubate for 1 h at 37°C. This was followed by a second 1 h 37°C extraction using a 1:1 mixture of 1 % formic acid in 50 mM TEAB and 100% acetonitrile extraction buffer. This secondary tryptic peptide extraction fraction was pooled with the aforementioned retained in-solution tryptic peptides. Pooled peptides were then dried, re-suspended, and desalted using ZipTip C18 pipette tips (ZTC18S960; Millipore), as previously described (Uhrig *et al*., 2019). Desalted peptides were then dried and re-suspended in 3% (v/v) ACN / 0.1% (v/v) formic acid immediately prior to MS analysis. Data was acquired using a Thermo Fisher Scientific Orbitrap Fusion Lumos Tribrid system. Peptides were eluted into the mass spectrometer using an nLC-1200 (Thermo Fisher Scientific) mounted with an ES901 column (Thermo Fisher Scientific). Peptides were eluted using a 66 min gradient of increasing Buffer B 0 – 18% (3 min), 18 – 28% (14 min), 28-46% (11 min), 46-100% (5 min); Buffer A: 3% ACN, 0.1 % FA; Buffer B: 80 % ACN, 0.1 % FA. All mass spectra were acquired in data dependent acquisition mode. MS1 data were acquired using a resolution of 30000, a scan range of 375-1700 m/z, a maximum injection time of 50 ms and a RF lens setting of 30%. MS2 data were acquired using the ion trap in rapid scan rate mode, with maximum injection time set to dynamic. A 30 % HCD collision energy was using for peptide fragmentation. Subsequently, all data was analyzed using MaxQuant 2.0.3.0 (Cox and Mann, 2008) using default parameters and the Araport 11 database (Cheng *et al*., 2017) with decoy mode set to revert. In brief, data search parameters included: trypsin cleavage permitting 2 missed cleavages, carbamidomethylation of cysteine residues (fixed modification), while methionine oxidation and phosphorylated serine/threonine/tyrosine were set as variable modifications. A PSM, peptide and protein FDR threshold of 0.01 was employed.

### Immunoblotting

Proteins were separated by 10% SDS-PAGE, transferred onto polyvinylidene difluoride membranes (BioRad), and blocked with 5% nonfat milk or bovine serum albumin (BSA) in Tris-buffered saline containing 0.05–0.1% Tween-20. MAZ-GFP and MAZ^K141N^-GFP accumulation blots were probed with mouse anti-GFP (1:5,000, Roche 1814460001) and goat anti-mouse-HRP (1:10,000, Sigma A0168). MAPK activation blots were probed with rabbit anti-p44/42 MAPK (Erk1/2) (1:2,000, Cell Signaling 9102S) and goat anti-rabbit-IgG (1:10,000, Sigma A0545). Membranes were incubated with ECL Clarity Substrate (BioRad), visualized on a ChemiDoc Touch Imaging System (BioRad) and subsequently stained with Coomassie Brilliant Blue (CBB) R-250 (MP Biomedicals) as a loading control.

### Statistics

GraphPad Prism 8 was used to perform statistical tests on all quantitative data, as indicated in figure legends.

## Data Availability

All raw proteomic data have been uploaded to ProteomeXchange (http://www.proteomexchange.org/) via the Proteomics IDEntification Database (PRIDE; https://www.ebi.ac.uk/pride/). Project Accession: PXD052679. Username; Password: hAT8rFqbBmdt

## Author Contributions

MGD and TD performed the majority of the work. MGD, TD, CGF, and JEM generated materials, performed experiments, and analyzed results. KRS and VNM generated materials. MCRG performed phosphoproteomics, supervised by RGU. MB and RD performed supporting experiments that are not shown, and RD curated phosphosites from online databases. Individual credits are included wherever possible in the figure captions and table legends. RS supervised JEM. JM designed the project, guided the work, supervised MGD, TD, CGF, MB, KRS, VNM, and RD, secured funding, and wrote the paper with input from all authors.

## Acknowledgements

We thank all members of the Monaghan Lab for their comments on this manuscript and for their commitment to fostering a welcoming and collaborative research environment. We thank Claire Wright for assistance with genotyping, Jack Moore for technical assistance and maintenance of the University of Alberta Mass Spectrometry and Proteomics Facility, Saeid Mobini for managing the Queen’s University Phytotron Facility, and the Center for Advanced Imaging (CAi) at Heinrich Heine University. Queen’s University is situated on the territory of the Haudenosaunee and Anishinaabek and we are grateful to live, work, and play on these lands.

## Supplemental Data

**Supplemental Table S1**. Primers, clones, and germplasm used in this study.

**Supplemental Figure S1.**
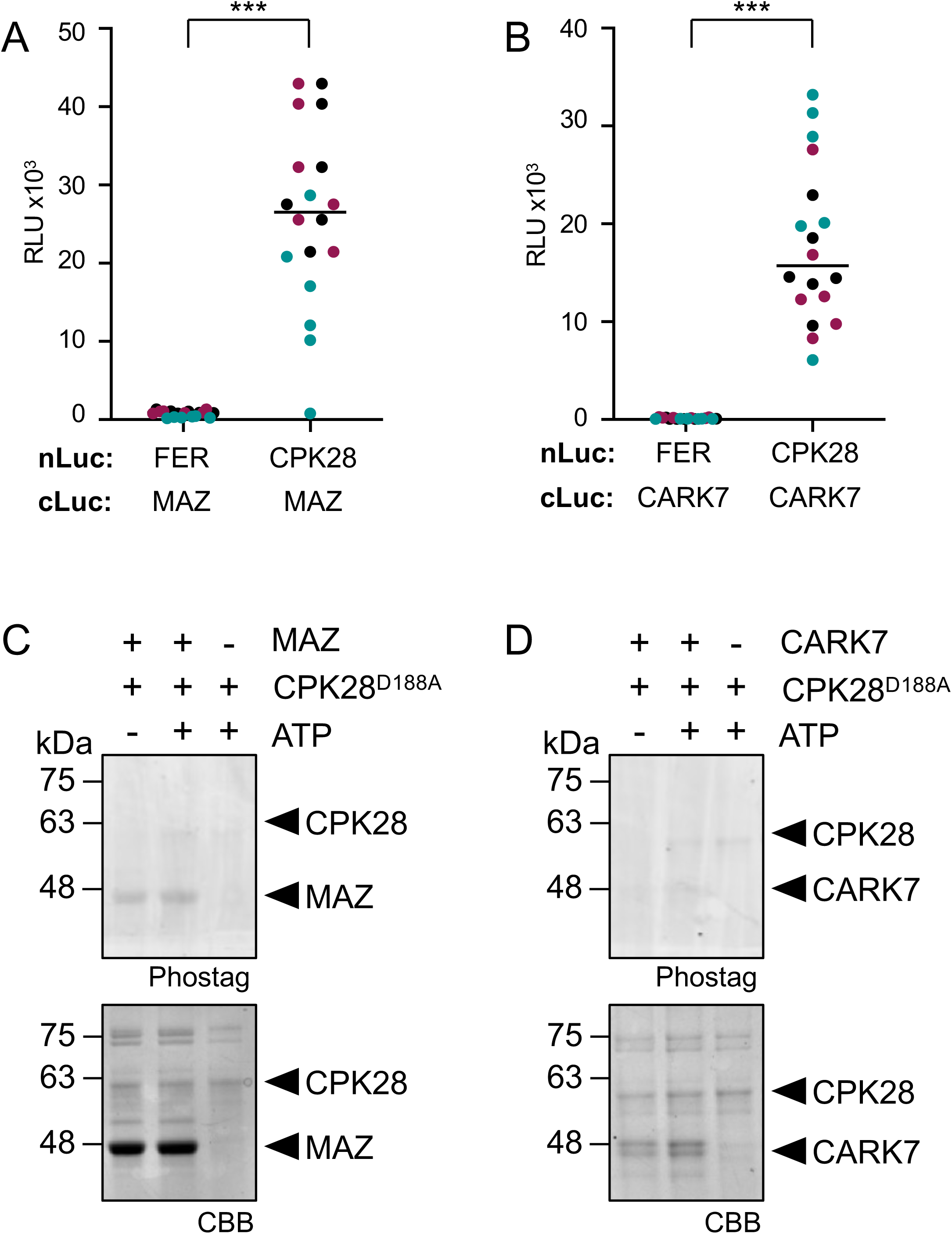
MAZ and CARK7 associate with but do not phosphorylate CPK28. **(A-B)** Split-luciferase complementation assays with FER-nLuc or CPK28-nLuc and cLuc-MAZ **(A)** or cLuc-CARK7 **(B)**. Total photon counts are plotted as relative light units (RLU) after co-expression of the respective proteins in *N. benthamiana*. Experiments were performed by CGF and MGD; data presented here were collected by MGD from 3 independent experiments (n=18), and are plotted together, denoted by black, teal, and magenta dots. The association between CPK28 and MAZ or CARK7 are significantly different from each control (Student’s unpaired t-test; p<0.0001). **(C-D)** *In vitro* kinase assays using His_6_-MAZ **(C)** or His_6_-CARK7 **(D)** as the kinase and catalytically-inactive His_6_-CPK28^D188A^ as the substrate, as visualized by Phostag staining. Protein loading is indicated by post-staining the membranes with Coomassie Brilliant Blue (CBB). Assays were performed more than 3 times each by TD with similar results; representative data are shown.

**Supplemental Figure S2.**
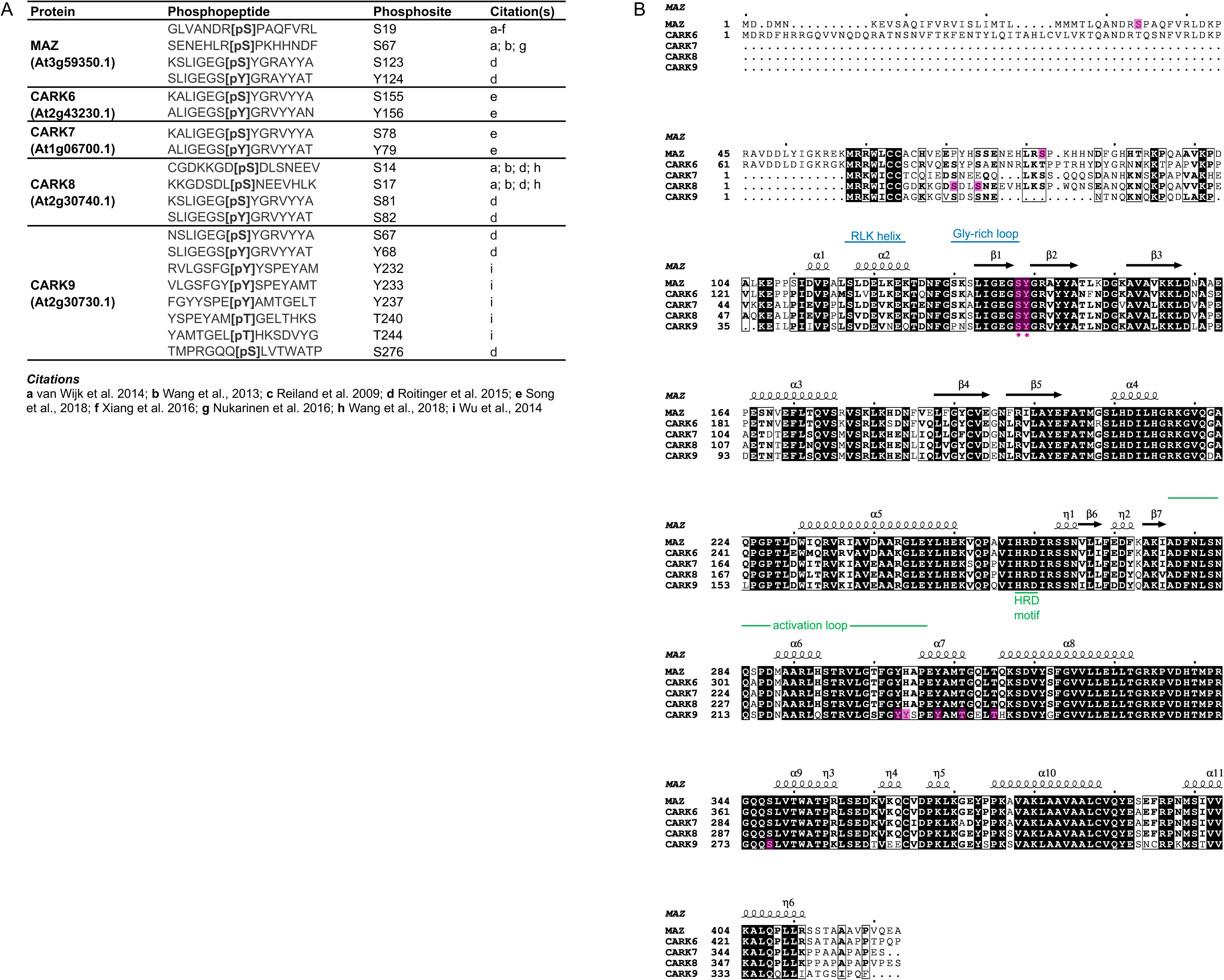
Phosphorylation sites identified on subgroup VIII-III RLCKs from various studies. **(A)** List of phosphorylation sites identified on subgroup VIII-III RLCKs from various phosphoproteomics studies, curated from online databases (Durek *et al*., 2010; Lin *et al*., 2021) based on the following publications (Reiland *et al*., 2009; Durek *et al*., 2010; Wang *et al*., 2013, 2018*b*; van Wijk *et al*., 2014; Wu *et al*., 2014; Roitinger *et al*., 2015; Nukarinen *et al*., 2016; Xiang *et al*., 2016; Song *et al*., 2018). **(B)** Multiple sequence alignment of subgroup VIII-III RLCKs with protein secondary structures indicated as visualized by ESPript (Robert and Gouet, 2014). Residues highlighted in magenta correspond to those in **(A)**; the magenta stars indicate highly conserved phosphorylation sites. Analysis done by RD.

**Supplemental Figure S3.**
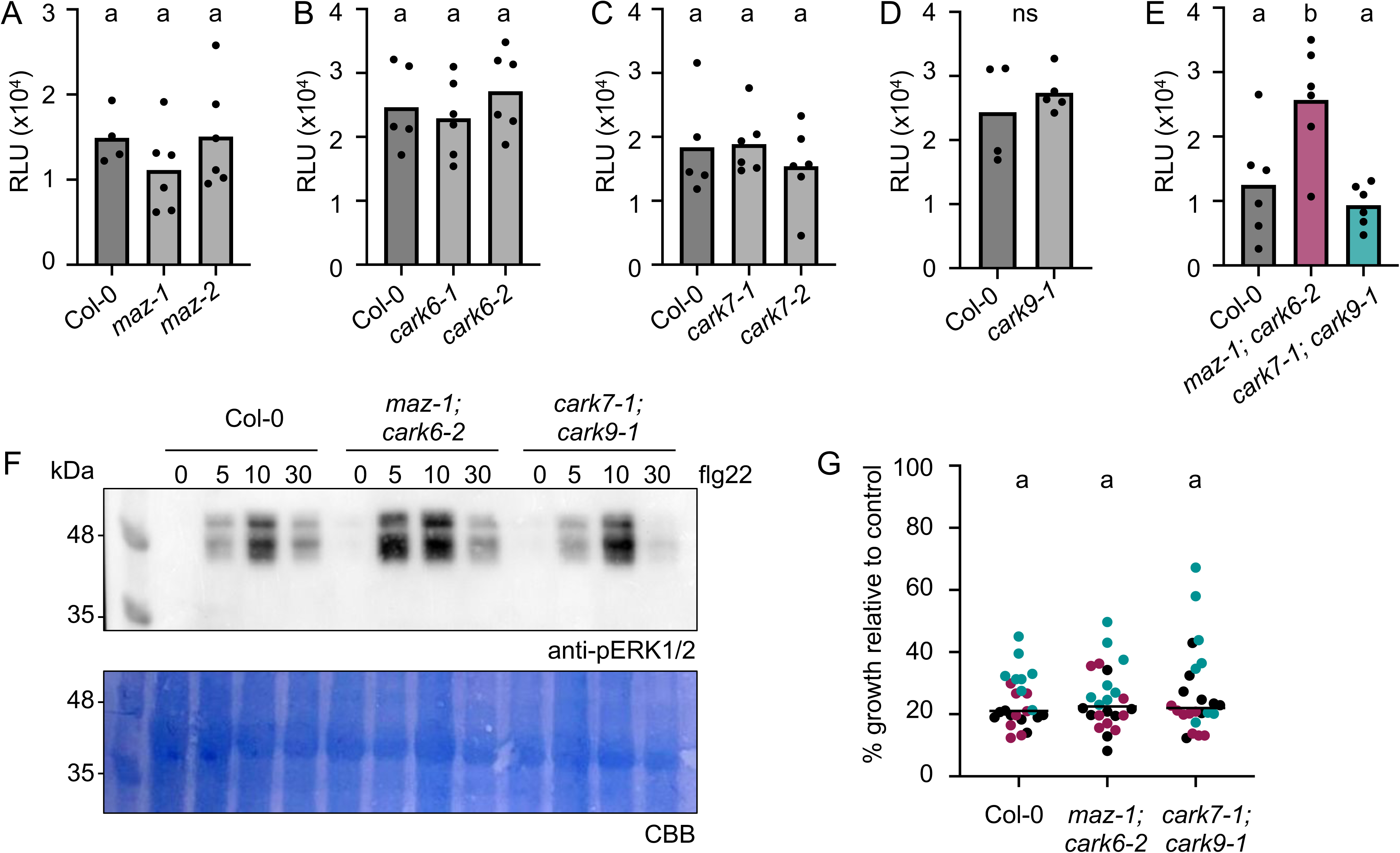
Analysis of immune responses in *maz-1; cark6-2* and *cark7-1; cark9-1* mutants. **(A-E)** ROS production measured in relative light units (RLUs) after treatment with 100 nM flg22. Values represent means +/-standard deviation (n=4-6). Data presented in **A-D** was collected by CGF; data in **E** was collected by MGD. Lower-case letters indicate statistically significant groups determined by a one-way ANOVA followed by Tukey’s post-hoc test (p=0.01). These assays were repeated several times over a 5 year period by CGF, MGD, and JM; representative experiments are shown. **(F)** Western blot indicating the activation of MAPKs before (0 min) and after exposure to 200 nM flg22 (5, 10, 30 min) in the indicated genotypes. The anti-pERK1/2 antibody recognizes the phosphorylated/activated forms of MPK6, MPK3, and MPK4/11. Coomassie Brilliant Blue (CBB) staining of the same membranes indicates loading. Experiments were completed 4 times with similar results by MGD; representative data is shown. **(G)** Growth inhibition of entire 12 day-old seedlings grown in liquid 0.5x MS media supplemented with 100 nM flg22, relative to mean seedling weight of 12 day-old seedlings grown in liquid 0.5x MS media with no peptide. Data from 3 independent biological replicates performed by MGD are plotted together, denoted by black, teal, and magenta dots. A one-way ANOVA followed by Tukey’s post-hoc test indicates no significant differences between groups, indicated by lower case letters. Credits for genetic crosses and genotyping are provided in **Supplemental Table S1**.

**Supplemental Figure S4.**
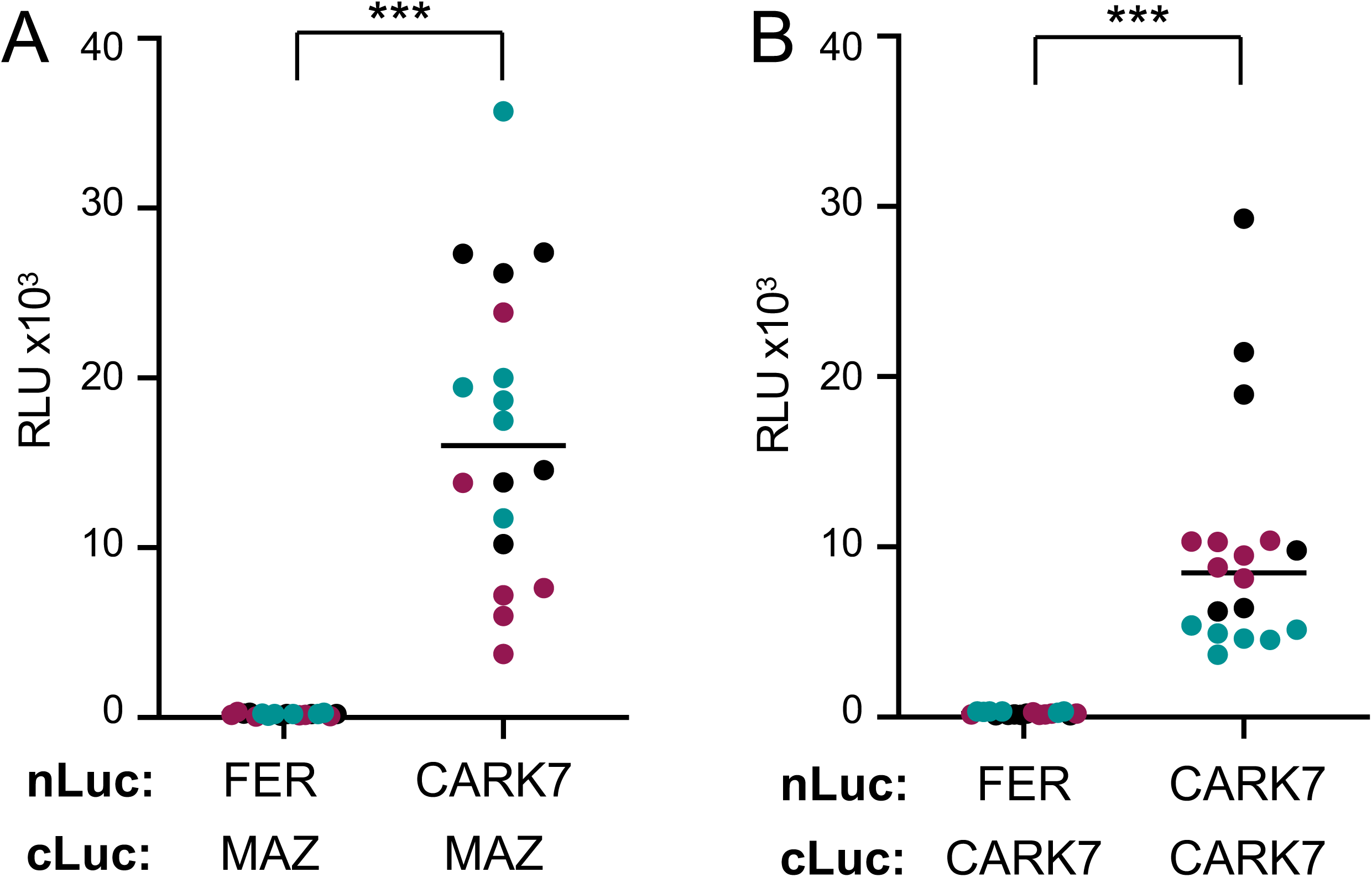
MAZ and CARK7 associate with themselves and each other. **(A-B)** Split-luciferase complementation assays with FER-nLuc or CARK7-nLuc and cLuc-MAZ **(A)** or cLuc-CARK7 **(B)**. Total photon counts are plotted as relative light units (RLU) after co-expression of the respective proteins in *N. benthamiana*. Experiments were performed by CGF and MGD; data presented here were collected by MGD from 3 independent experiments (n=18), and are plotted together, denoted by black, teal, and magenta dots. The association between CARK7-MAZ and CARK7-CARK7 are significantly different from the controls (Student’s unpaired t-test; p<0.0001).

**Supplemental Figure S5.**
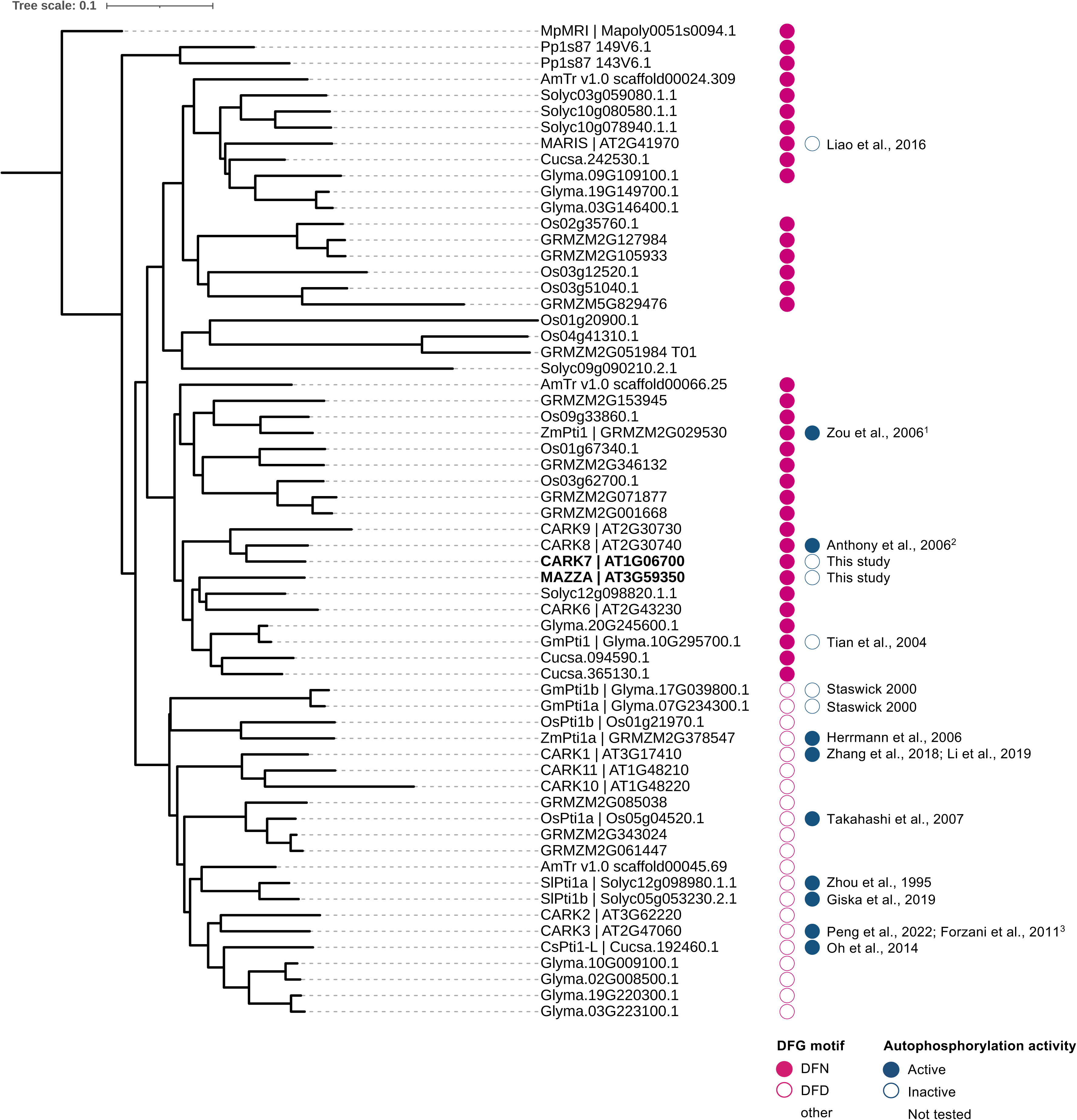
Analysis of the DFG motif in subgroup VIII RLCKs from multiple species. Phylogeny of subgroup VIII RLCKs from *Amborella trichopoda* (3 members), *Arabidopsis thaliana* (11 members), *Solanum lycopersicum* (7 members), *Glycine max* (11 members), *Cucumis sativus* (3 members), *Oryza sativa* (10 members), *Zea mays* (13 members), *Physcomitrium patens* (2 members), and *Marchantia polymorpha* (1 member). A multiple sequence alignment was generated using MUSCLE in MEGAX (Kumar *et al*., 2018) and used to generate a neighbour-joining tree, visualized in iTOL (Letunic and Bork, 2021). Proteins containing ‘DFN’ or ‘DFD’ motifs in place of the canonical ‘DFG’ motif are indicated by closed and open pink circles, respectively, while proteins containing motifs other than DFN or DFD are indicated by blank spaces. Proteins that have demonstrated autophosphorylation activity are indicated by closed blue circles, while those that are inactive are indicated by open blue circles. Proteins that have not yet been tested are indicated by blank spaces. Analysis done by JM.

## References

1. Anthony RG, Khan S, Costa J, Pais MS, Bögre L. 2006. The Arabidopsis Protein Kinase PTI1-2 Is Activated by Convergent Phosphatidic Acid and Oxidative Stress Signaling Pathways Downstream of PDK1 and OXI1*. The Journal of biological chemistry 281, 37536–37546.

2. Asai T, Tena G, Plotnikova J, Willmann MR, Chiu W-L, Gomez-Gomez L, Boller T, Ausubel FM, Sheen J. 2002. MAP kinase signalling cascade in Arabidopsis innate immunity. Nature 415, 977–983.

3. Bender KW, Couto D, Kadota Y, et al. 2021. Activation loop phosphorylation of a non-RD receptor kinase initiates plant innate immune signaling. Proceedings of the National Academy of Sciences of the United States of America 118 (38) e2108242118.

4. Bleckmann A, Weidtkamp-Peters S, Seidel CAM, Simon R. 2010. Stem cell signaling in Arabidopsis requires CRN to localize CLV2 to the plasma membrane. Plant physiology 152, 166–176.

5. Blümke P, Schlegel J, Gonzalez-Ferrer C, Becher S, Pinto KG, Monaghan J, Simon R. 2021. Receptor-like cytoplasmic kinase MAZZA mediates developmental processes with CLAVATA1 family receptors in Arabidopsis. Journal of experimental botany 72, 4853–4870.

6. Boisson-Dernier A, Franck CM, Lituiev DS, Grossniklaus U. 2015. Receptor-like cytoplasmic kinase MARIS functions downstream of CrRLK1L-dependent signaling during tip growth. Proceedings of the National Academy of Sciences of the United States of America 112, 12211–12216.

7. Bredow M, Bender KW, Johnson Dingee A, et al. 2021. Phosphorylation-dependent subfunctionalization of the calcium-dependent protein kinase CPK28. Proceedings of the National Academy of Sciences of the United States of America 118 (19) e2024272118.

8. Bredow M, Sementchoukova I, Siegel K, Monaghan J. 2019. Pattern-Triggered Oxidative Burst and Seedling Growth Inhibition Assays in Arabidopsis thaliana. Journal of visualized experiments: JoVE doi: 10.3791/59437.

9. Chandra S, Martin GB, Low PS. 1996. The Pto kinase mediates a signaling pathway leading to the oxidative burst in tomato. Proceedings of the National Academy of Sciences of the United States of America 93, 13393–13397.

10. Cheng C-Y, Krishnakumar V, Chan AP, Thibaud-Nissen F, Schobel S, Town CD. 2017. Araport11: a complete reannotation of the Arabidopsis thaliana reference genome. The Plant journal: for cell and molecular biology 89, 789–804.

11. Chen H, Zou Y, Shang Y, Lin H, Wang Y, Cai R, Tang X, Zhou J-M. 2008. Firefly luciferase complementation imaging assay for protein-protein interactions in plants. Plant physiology 146, 368–376.

12. Chinchilla D, Zipfel C, Robatzek S, Kemmerling B, Nürnberger T, Jones JDG, Felix G, Boller T. 2007. A flagellin-induced complex of the receptor FLS2 and BAK1 initiates plant defence. Nature 448, 497–500.

13. Clough SJ, Bent AF. 1998. Floral dip: a simplified method for Agrobacterium-mediated transformation of Arabidopsis thaliana. The Plant journal: for cell and molecular biology 16, 735–743.

14. Cox J, Mann M. 2008. MaxQuant enables high peptide identification rates, individualized p.p.b.-range mass accuracies and proteome-wide protein quantification. Nature biotechnology 26, 1367–1372.

15. Cutler SR, Ehrhardt DW, Griffitts JS, Somerville CR. 2000. Random GFP::cDNA fusions enable visualization of subcellular structures in cells of Arabidopsis at a high frequency. Proceedings of the National Academy of Sciences of the United States of America 97, 3718–3723.

16. Dardick C, Ronald P. 2006. Plant and animal pathogen recognition receptors signal through non-RD kinases. PLoS pathogens 2, e2.

17. DeFalco TA, Zipfel C. 2021. Molecular mechanisms of early plant pattern-triggered immune signaling. Molecular cell 81, 4346.

18. Dias MG, Soleimani F, Monaghan J. 2022. Activation and turnover of the plant immune signaling kinase BIK1: a fine balance. Essays in biochemistry 66, 207–218.

19. Dias MG, Doss B, Rawat A, et al. 2023. Subfamily C7 Raf-like kinases MRK1, RAF26, and RAF39 regulate immune homeostasis and stomatal opening in Arabidopsis thaliana. bioRxiv, 2023.11.29.569073.

20. Ding Y, Yang H, Wu S, Fu D, Li M, Gong Z, Yang S. 2022. CPK28-NLP7 module integrates cold-induced Ca^2+^ signal and transcriptional reprogramming in *Arabidopsis*. Science advances 8, eabn7901.

21. Durek P, Schmidt R, Heazlewood JL, Jones A, MacLean D, Nagel A, Kersten B, Schulze WX. 2010. PhosPhAt: the Arabidopsis thaliana phosphorylation site database. An update. Nucleic acids research 38, D828–34.

22. Forzani C, Carreri A, de la Fuente van Bentem S, Lecourieux D, Lecourieux F, Hirt H. 2011. The Arabidopsis protein kinase Pto-interacting 1-4 is a common target of the oxidative signal-inducible 1 and mitogen-activated protein kinases. The FEBS journal 278, 1126–1136.

23. Giska F, Martin GB. 2019. PP2C phosphatase Pic1 negatively regulates the phosphorylation status of Pti1b kinase, a regulator of flagellin-triggered immunity in tomato. Biochemical Journal 476, 1621–1635.

24. Gómez-Gómez L, Felix G, Boller T. 1999. A single locus determines sensitivity to bacterial flagellin in Arabidopsis thaliana. The Plant journal: for cell and molecular biology 18, 277–284.

25. Heese A, Hann DR, Gimenez-Ibanez S, Jones AME, He K, Li J, Schroeder JI, Peck SC, Rathjen JP. 2007. The receptor-like kinase SERK3/BAK1 is a central regulator of innate immunity in plants. Proceedings of the National Academy of Sciences of the United States of America 104, 12217–12222.

26. Herrmann MM, Pinto S, Kluth J, Wienand U, Lorbiecke R. 2006. The PTI1-like kinase ZmPti1a from maize (Zea mays L.) co-localizes with callose at the plasma membrane of pollen and facilitates a competitive advantage to the male gametophyte. BMC plant biology 6, 22.

27. Holmes DR, Bredow M, Thor K, Pascetta SA, Sementchoukova I, Siegel KR, Zipfel C, Monaghan J. 2021. A novel allele of the Arabidopsis thaliana MACPF protein CAD1 results in deregulated immune signaling. Genetics 217 (4) iyab022.

28. Huangfu Y, Pan J, Li Z, Wang Q, Mastouri F, Li Y, Yang S, Liu M, Dai S, Liu W. 2021. Genome-wide identification of PTI1 family in Setaria italica and salinity-responsive functional analysis of SiPTI1-5. BMC plant biology 21, 319.

29. Jia Y, Loh YT, Zhou J, Martin GB. 1997. Alleles of Pto and Fen occur in bacterial speck-susceptible and fenthion-insensitive tomato cultivars and encode active protein kinases. The Plant cell 9, 61–73.

30. Jumper J, Evans R, Pritzel A, et al. 2021. Highly accurate protein structure prediction with AlphaFold. Nature 596, 583–589.

31. Karimi M, Inzé D, Depicker A. 2002. GATEWAY vectors for Agrobacterium-mediated plant transformation. Trends in plant science 7, 193–195.

32. Kondo T, Sawa S, Kinoshita A, Mizuno S, Kakimoto T, Fukuda H, Sakagami Y. 2006. A plant peptide encoded by CLV3 identified by in situ MALDI-TOF MS analysis. Science 313, 845–848.

33. Kourelis J. 2023. Interplay between cell-surface receptor and intracellular NLR-mediated immune responses. The New phytologist 240, 2218–2226.

34. Kumar S, Stecher G, Li M, Knyaz C, Tamura K. 2018. MEGA X: Molecular Evolutionary Genetics Analysis across Computing Platforms. Molecular biology and evolution 35, 1547–1549.

35. Kung JE, Jura N. 2016. Structural Basis for the Non-catalytic Functions of Protein Kinases. Structure 24, 7–24.

36. Lehti-Shiu MD, Shiu S-H. 2012. Diversity, classification and function of the plant protein kinase superfamily. Philosophical transactions of the Royal Society of London. Series B, Biological sciences 367, 2619–2639.

37. Letunic I, Bork P. 2021. Interactive Tree Of Life (iTOL) v5: an online tool for phylogenetic tree display and annotation. Nucleic acids research 49, W293–W296.

38. Liang X, Zhou J-M. 2018. Receptor-Like Cytoplasmic Kinases: Central Players in Plant Receptor Kinase–Mediated Signaling. Annual review of plant biology 69, 267–299.

39. Liao H-Z, Zhu M-M, Cui H-H, Du X-Y, Tang Y, Chen L-Q, Ye D, Zhang X-Q. 2016. MARIS plays important roles in Arabidopsis pollen tube and root hair growth. Journal of integrative plant biology 58, 927–940.

40. Li X, Kong X, Huang Q, et al. 2019. CARK1 phosphorylates subfamily III members of ABA receptors. Journal of experimental botany 70, 519–528.

41. Lin S, Wang C, Zhou J, Shi Y, Ruan C, Tu Y, Yao L, Peng D, Xue Y. 2021. EPSD: a well-annotated data resource of protein phosphorylation sites in eukaryotes. Briefings in bioinformatics 22, 298–307.

42. Li X, Xie Y, Zhang Q, et al. 2022. Monomerization of abscisic acid receptors through CARKs-mediated phosphorylation. The New phytologist 235, 533–549.

43. Lu D, Wu S, Gao X, Zhang Y, Shan L, He P. 2010. A receptor-like cytoplasmic kinase, BIK1, associates with a flagellin receptor complex to initiate plant innate immunity. Proceedings of the National Academy of Sciences of the United States of America 107, 496–501.

44. Martel A, Laflamme B, Seto D, Bastedo DP, Dillon MM, Almeida RND, Guttman DS, Desveaux D. 2020. Immunodiversity of the Arabidopsis ZAR1 NLR Is Conveyed by Receptor-Like Cytoplasmic Kinase Sensors. Frontiers in plant science 11, 1290.

45. Matschi S, Werner S, Schulze WX, Legen J, Hilger HH, Romeis T. 2013. Function of calcium-dependent protein kinase CPK28 of Arabidopsis thaliana in plant stem elongation and vascular development. The Plant journal: for cell and molecular biology 73, 883–896.

46. Matsui H, Yamazaki M, Kishi-Kaboshi M, Takahashi A, Hirochika H. 2010. AGC kinase OsOxi1 positively regulates basal resistance through suppression of OsPti1a-mediated negative regulation. Plant & cell physiology 51, 1731–1744.

47. Monaghan J, Matschi S, Shorinola O, et al. 2014. The calcium-dependent protein kinase CPK28 buffers plant immunity and regulates BIK1 turnover. Cell host & microbe 16, 605–615.

48. Monaghan J, Zipfel C. 2012. Plant pattern recognition receptor complexes at the plasma membrane. Current opinion in plant biology 15, 349–357.

49. Mucyn TS, Clemente A, Andriotis VME, Balmuth AL, Oldroyd GED, Staskawicz BJ, Rathjen JP. 2006. The tomato NBARC-LRR protein Prf interacts with Pto kinase in vivo to regulate specific plant immunity. The Plant cell 18, 2792–2806.

50. Mucyn TS, Wu A-J, Balmuth AL, Arasteh JM, Rathjen JP. 2009. Regulation of tomato Prf by Pto-like protein kinases. Molecular plant-microbe interactions: MPMI 22, 391–401.

51. Mühlenbeck H, Tsutsui Y, Lemmon MA, Bender KW, Zipfel C. 2024. Allosteric activation of the co-receptor BAK1 by the EFR receptor kinase initiates immune signaling. bioRxiv: the preprint server for biology doi: 10.1101/2023.08.23.554490.

52. Nam KH, Li J. 2002. BRI1/BAK1, a receptor kinase pair mediating brassinosteroid signaling. Cell 110, 203–212.

53. Nolen B, Taylor S, Ghosh G. 2004. Regulation of protein kinases; controlling activity through activation segment conformation. Molecular cell 15, 661–675.

54. Nukarinen E, Nägele T, Pedrotti L, et al. 2016. Quantitative phosphoproteomics reveals the role of the AMPK plant ortholog SnRK1 as a metabolic master regulator under energy deprivation. Scientific reports 6, 31697.

55. Oh S-K, Jang HA, Lee SS, Cho HS, Lee D-H, Choi D, Kwon S-Y. 2014. Cucumber Pti1-L is a cytoplasmic protein kinase involved in defense responses and salt tolerance. Journal of plant physiology 171, 817–822.

56. Peng L, He J, Yao H, et al. 2022. CARK3-mediated ADF4 regulates hypocotyl elongation and soil drought stress in Arabidopsis. Frontiers in plant science 13, 1065677.

57. Pettersen EF, Goddard TD, Huang CC, Meng EC, Couch GS, Croll TI, Morris JH, Ferrin TE. 2021. UCSF ChimeraX: Structure visualization for researchers, educators, and developers. Protein science: a publication of the Protein Society 30, 70–82.

58. Rao S, Zhou Z, Miao P, Bi G, Hu M, Wu Y, Feng F, Zhang X, Zhou J-M. 2018. Roles of Receptor-Like Cytoplasmic Kinase VII Members in Pattern-Triggered Immune Signaling. Plant physiology 177, 1679–1690.

59. Reiland S, Messerli G, Baerenfaller K, Gerrits B, Endler A, Grossmann J, Gruissem W, Baginsky S. 2009. Large-scale Arabidopsis phosphoproteome profiling reveals novel chloroplast kinase substrates and phosphorylation networks. Plant physiology 150, 889–903.

60. Rentel MC, Lecourieux D, Ouaked F, et al. 2004. OXI1 kinase is necessary for oxidative burst-mediated signalling in Arabidopsis. Nature 427, 858–861.

61. Robert X, Gouet P. 2014. Deciphering key features in protein structures with the new ENDscript server. Nucleic acids research 42, W320–4.

62. Roitinger E, Hofer M, Köcher T, Pichler P, Novatchkova M, Yang J, Schlögelhofer P, Mechtler K. 2015. Quantitative phosphoproteomics of the ataxia telangiectasia-mutated (ATM) and ataxia telangiectasia-mutated and rad3-related (ATR) dependent DNA damage response in Arabidopsis thaliana. Molecular & cellular proteomics: MCP 14, 556–571.

63. Schwizer S, Kraus CM, Dunham DM, Zheng Y, Fernandez-Pozo N, Pombo MA, Fei Z, Chakravarthy S, Martin GB. 2017. The tomato kinase Pti1 contributes to production of reactive oxygen species in response to two flagellin-derived peptides and promotes resistance to Pseudomonas syringae infection. Molecular plant-microbe interactions: MPMI 30, 725–738.

64. Segonzac C, Monaghan J. 2019. Modulation of plant innate immune signaling by small peptides. Current opinion in plant biology 51, 22–28.

65. Shirasu K. 2009. The HSP90-SGT1 chaperone complex for NLR immune sensors. Annual review of plant biology 60, 139–164.

66. Shiu SH, Bleecker AB. 2001. Receptor-like kinases from Arabidopsis form a monophyletic gene family related to animal receptor kinases. Proceedings of the National Academy of Sciences of the United States of America 98, 10763–10768.

67. Shu L-J, Kahlon PS, Ranf S. 2023. The power of patterns: new insights into pattern-triggered immunity. The New phytologist 240, 960–967.

68. Son Y, Cheong Y-K, Kim N-H, Chung H-T, Kang DG, Pae H-O. 2011. Mitogen-Activated Protein Kinases and Reactive Oxygen Species: How Can ROS Activate MAPK Pathways? Journal of signal transduction 2011, 792639.

69. Song G, Brachova L, Nikolau BJ, Jones AM, Walley JW. 2018. Heterotrimeric G-Protein-Dependent Proteome and Phosphoproteome in Unstimulated Arabidopsis Roots. Proteomics 18, e1800323.

70. Spatola Rossi T, Pain C, Botchway SW, Kriechbaumer V. 2022. FRET-FLIM to Determine Protein Interactions and Membrane Topology of Enzyme Complexes. Current protocols 2, e598.

71. Staswick P. 2000. Two expressed soybean genes with high sequence identity to tomato Pti1 kinase lack autophosphorylation activity. Archives of biochemistry and biophysics 383, 233–237.

72. Takahashi A, Agrawal GK, Yamazaki M, Onosato K, Miyao A, Kawasaki T, Shimamoto K, Hirochika H. 2007. Rice Pti1a negatively regulates RAR1-dependent defense responses. The Plant cell 19, 2940–2951.

73. Tang D, Wang G, Zhou J-M. 2017. Receptor Kinases in Plant-Pathogen Interactions: More Than Pattern Recognition. The Plant cell 29, 618–637.

74. Tian A-G, Luo G-Z, Wang Y-J, Zhang J-S, Gai J-Y, Chen S-Y. 2004. Isolation and characterization of a Pti1 homologue from soybean. Journal of experimental botany 55, 535–537.

75. Uhrig RG, Schläpfer P, Roschitzki B, Hirsch-Hoffmann M, Gruissem W. 2019. Diurnal changes in concerted plant protein phosphorylation and acetylation in Arabidopsis organs and seedlings. The Plant journal: for cell and molecular biology 99, 176–194.

76. Ung PM-U, Schlessinger A. 2015. DFGmodel: predicting protein kinase structures in inactive states for structure-based discovery of type-II inhibitors. ACS chemical biology 10, 269–278.

77. Veronese P, Nakagami H, Bluhm B, Abuqamar S, Chen X, Salmeron J, Dietrich RA, Hirt H, Mengiste T. 2006. The membrane-anchored BOTRYTIS-INDUCED KINASE1 plays distinct roles in Arabidopsis resistance to necrotrophic and biotrophic pathogens. The Plant cell 18, 257–273.

78. Voinnet O, Rivas S, Mestre P, Baulcombe D. 2003. An enhanced transient expression system in plants based on suppression of gene silencing by the p19 protein of tomato bushy stunt virus. The Plant journal: for cell and molecular biology 33, 949– 956.

79. Wang X, Bian Y, Cheng K, Gu L-F, Ye M, Zou H, Sun SS-M, He J-X. 2013. A large-scale protein phosphorylation analysis reveals novel phosphorylation motifs and phosphoregulatory networks in Arabidopsis. Journal of proteomics 78, 486–498.

80. Wang J, Grubb LE, Wang J, et al. 2018*a*. A Regulatory Module Controlling Homeostasis of a Plant Immune Kinase. Molecular cell 69, 493–504.e6.

81. Wang G, Roux B, Feng F, et al. 2015. The Decoy Substrate of a Pathogen Effector and a Pseudokinase Specify Pathogen-Induced Modified-Self Recognition and Immunity in Plants. Cell host & microbe 18, 285–295.

82. Wang K, Yang Z, Qing D, et al. 2018*b*. Quantitative and functional posttranslational modification proteomics reveals that TREPH1 plays a role in plant touch-delayed bolting. Proceedings of the National Academy of Sciences of the United States of America 115, E10265–E10274.

83. van Wijk KJ, Friso G, Walther D, Schulze WX. 2014. Meta-Analysis of Arabidopsis thaliana Phospho-Proteomics Data Reveals Compartmentalization of Phosphorylation Motifs. The Plant cell 26, 2367–2389.

84. Wu X, Sklodowski K, Encke B, Schulze WX. 2014. A kinase-phosphatase signaling module with BSK8 and BSL2 involved in regulation of sucrose-phosphate synthase. Journal of proteome research 13, 3397–3409.

85. Xiang Y, Song B, Née G, Kramer K, Finkemeier I, Soppe WJJ. 2016. Sequence Polymorphisms at the REDUCED DORMANCY5 Pseudophosphatase Underlie Natural Variation in Arabidopsis Dormancy. Plant physiology 171, 2659–2670.

86. Yu X, Feng B, He P, Shan L. 2017. From Chaos to Harmony: Responses and Signaling upon Microbial Pattern Recognition. Annual review of phytopathology 55, 109–137.

87. Zhang L, Li X, Li D, et al. 2018. CARK1 mediates ABA signaling by phosphorylation of ABA receptors. Cell discovery 4, 30.

88. Zhang J, Li W, Xiang T, et al. 2010. Receptor-like Cytoplasmic Kinases Integrate Signaling from Multiple Plant Immune Receptors and Are Targeted by a Pseudomonas syringae Effector. Cell host & microbe 7, 290–301.

89. Zhou J, Loh YT, Bressan RA, Martin GB. 1995. The tomato gene Pti1 encodes a serine/threonine kinase that is phosphorylated by Pto and is involved in the hypersensitive response. Cell 83, 925–935.

90. Zou H, Wu Z, Yang Q, Zhang X, Cao M, Jia W, Huang C, Xiao X. 2006. Gene expression analyses of ZmPti1, encoding a maize Pti-like kinase, suggest a role in stress signaling. Plant science: an international journal of experimental plant biology 171, 99–105.

